# Ectopic expression induces abnormal somatodendritic distribution of tau in the mouse brain

**DOI:** 10.1101/457853

**Authors:** Atsuko Kubo, Shouyou Ueda, Ayaka Yamane, Satoko Wada-Kakuda, Makoto Matsuyama, Akane Nomori, Akihiko Takashima, Motohito Goto, Mamoru Ito, Takami Tomiyama, Hiroshi Mori, Yasuo Ihara, Hiroaki Misonou, Tomohiro Miyasaka

## Abstract

Tau is a microtubule-associated protein that is localized to the axon. In Alzheimer’s disease, the distribution of tau undergoes a remarkable alteration, leading to the formation of tau inclusions in the somatodendritic compartment. To investigate how this mis-localization occurs, we recently developed immunohistochemical techniques that can separately detect endogenous mouse and exogenous human tau with high sensitivity, which allows us to visualize not only the pathological but pre-aggregated tau in mouse brain tissues of both sex. In tau transgenic mouse brains, exogenous human tau was abundant in dendrites and somata even in the presymptomatic period, whereas the axonal localization of endogenous mouse tau was unaffected. In stark contrast, exogenous tau was properly localized to the axon in human tau knock-in mice. We tracked this difference to the temporal expression patterns of tau. Tau mRNA was continuously expressed in the transgenic mice, whereas endogenous tau and exogenous tau in the knock-in mice exhibited high expression levels during the neonatal period and strong suppression into the adulthood. These results indicated the uncontrolled expression of exogenous tau beyond the developmental period as a cause of mis-localization in the transgenic mice. Super-resolution microscopic and biochemical analyses also indicated that the interaction between microtubules and exogenous tau was indeed impaired in the tau transgenic mice. Thus, the ectopic expression of tau may be critical for its somatodendritic mis-localization, a key step of the tauopathy.

**Significance Statement:** Somatodendritic localization of tau may be an early step leading to the neuronal degeneration in tauopathies. However, the mechanisms of the normal axonal distribution of tau and the mis-localization of pathological tau remain obscure. Our immunohistochemical and biochemical analyses demonstrated that the endogenous mouse tau is transiently expressed in neonatal brains, that exogenous human tau expressed corresponding to such tau expression profile can distribute into the axon, and that the constitutive expression of tau into adulthood (ex. human tau in Tg mice) results in abnormal somatodendritic localization. Thus, the expression profile of tau is tightly associated with the localization of tau, and the ectopic expression of tau in matured neurons may be involved in the pathogenesis of tauopathy.

## Introduction

Tau is a microtubule (MT)-associated protein that is preferentially expressed in neuronal cells; within neurons, tau is exclusively expressed in axons (Scholz and Mandelkow, 2014). Tau is also known as a component of the paired helical filament that is found in neurofibrillary tangles (NFTs) or neuropil threads in “tauopathies”, including Alzheimer’s disease (AD) (Johnson and Jenkins, 1999). Both pathological evidence, which indicates a strong correlation between the formation of tau pathologies and neuronal degeneration (Gomez-Isla et al., 1997; Delacourte et al., 1999), and genetic evidence strongly suggest that tau can directly cause neurodegeneration and dementia (Ghetti et al., 2015).

Despite the axonal localization of tau in normal neurons (Kubo et al., in press), in AD and other tauopathies, tau inclusions are formed in the somatodendritic compartments of affected neurons (Kowall and Kosik, 1987; Braak and Braak, 1994). Cumulative evidence indicates that the formation of NFTs itself might not directly cause neuronal dysfunction and degeneration (Miyasaka et al., 2005a; Santacruz et al., 2005; Kuchibhotla et al., 2014) and that the abnormal distribution of presumably unaggregated tau into dendrites or spines is a critical determinant for neurodegeneration (Zempel et al., 2010; Frandemiche et al., 2014). Therefore, the "mis-localization" of tau may be a key step in the pathogenesis of tauopathies (Zempel and Mandelkow, 2014).

To determine how axonal tau mis-localizes to and accumulates in the somatodendritic compartment, we analyzed the distribution of endogenous tau and compared it with that of exogenously expressed human tau in common mouse models of tauopathy, which exhibit tau pathologies. Although several previous studies have shown the overall distribution of tau in normal brain tissues (Binder et al., 1985; Kowall and Kosik, 1987; Viereck et al., 1988; Trojanowski et al., 1989), the precise subcellular localization of endogenous tau and how this localization pattern changes in tauopathy models have not yet been extensively demonstrated, presumably due to the poor antigenicity of unaggregated endogenous tau (Trojanowski et al., 1989). We have recently developed new antibodies against tau and optimized procedures to reliably detect endogenous normal, unaggregated tau and expressed human tau separately in brain tissues (Kubo et al., in press). Here, using these new techniques, we investigated how its localization is affected in the animal models of tauopathy.

## Materials and Methods

### Animals and human tissues

Following animals were used in this study; P301L tau-transgenic mice (P301L-Tg; (Kimura et al., 2010)), Wild-type tau transgenic mice (WTtau-Tg; (Kimura et al., 2007)), tau knockout mice (tau-KO; (Dawson et al., 2001)), Line 264-Tg tau mice (tau264; (Umeda et al., 2013)), Thy1-EGFP mouse (M line; (Feng et al., 2000)), and PS19 mice. PS19 mice, which express P301S tau under the mouse prion promoter, were purchased from The Jackson Laboratory (Yoshiyama et al., 2007). P301L tau knock-in mice (P301L-KI) were developed by inserting human cDNA and floxed PGK-Neo poly-A cassette in the same region as the previous tau knockout mice (Harada et al., 1994). Namely, the knock-in mice were developed by in-frame insertion of human cDNA for 0N4R isoform (383 amino acids) followed by rabbit β-globin poly-A signal sequence from the mouse endogenous start codon in the first coding exon. Floxed PGK neo poly-A was inserted after the human tau cDNA (Matsumura et al., 1999). All animal experiments were approved by the institutional animal care and use committees. Both male and female animals were used.

### Antibodies

Anti-tauN and anti-MAP2N rabbit polyclonal antibodies were raised against the N-term peptide (AEPRQEFEVMEDHAGGGC) of human tau and the N-terminal 150 amino acid fragment of recombinant human MAP2, respectively. Anti-human tau peptide rabbit polyclonal antibody was raised against the peptide (GTYGLGDRKDQGGYTMHQGGC). Anti-rodent tauN (anti-RtauN) rabbit polyclonal antibody was raised against the peptide (DTMEDHAGDYTLLQDEG) corresponding to the N-terminal potion of mouse tau. Anti-total tau (RTM38), anti-human tau specific (RTM49) and anti-mouse tau specific (RTM47) rat monoclonal antibodies were raised against purified recombinant human and mouse tau, respectively. We have reported the production and validation of these antibodies recently (Kubo et al., in press). The anti-tau rat IgGs are now commercially available from FUJIFILM Wako Pure Chemical Corporation (Osaka, Japan)).

### Antibody dilutions

Dilutions of the antibodies for immunostaining are listed below; anti-tauN, anti-MAP2N, and anti-RtauN (1:1000), anti-tau12 (tau12, mouse monoclonal antibody, 1:2000, Abcam, Cambridge, UK), tau1 (tau1, mouse monoclonal antibody, 1:1000, Millipore, Billerica, MA), anti-MAP2 (HM2, mouse monoclonal antibody, 1:1000, Sigma-Aldrich, St. Louis, MO), RTM49, RTM47 and RTM38 (1:300), tau5 (tau5, mouse monoclonal antibody, 1:1000, Millipore), BD-anti-tau (anti-tau, mouse monoclonal antibody, 1:1000, Becton, Dickinson and Company, Franklin Lakes, NJ), anti-NeuN (mouse monoclonal antibody, 1:500, Abcam), anti-S100b (rabbit monoclonal antibody, 1:500, Novus Biologicals, Inc., Littleton, CO), Iba-1 (rabbit polyclonal, 1:1000, Wako Pure Chemical Industries, Osaka, Japan), anti-olig2 (rabbit polyclonal, 1:500, Proteintech group, Rosemont, IL), anti-Drebrin (Drebrin, mouse monoclonal antibody, 1:1000, Novus Biological, Littleton, CO), anti-VGlut1 (VGlut1, guinea pig polyclonal antibody, 1:1000, Synaptic Systems, Goettingen, Germany), anti-Pan-Axonal Neurofilament maker (SMI-312R, mouse monoclonal antibody, 1:1000, Covance, Princeton, NJ), anti-phosphorylated neurofilament (SMI31, 1:500, Covance), anti-amyloid β (anti-Human amyloid β(N), rabbit polyclonal antibody, 1:500, IBL, Takasaki, Japan), AT8 (mouse monoclonal antibody, 1:200, Innogenetics, Ghent, Belgium), anti-MBP (rat monoclonal antibody, 1:200, Abcam,) and Alexa 488-conjugated DM1A (DM1A-488, 1:100, Millipore).

### Preparation of tissue sections and neuronal primary cultures

Animals were anesthetized with pentobarbital and fixed via transcardial perfusion with 4% paraformaldehyde (PFA) in phosphate buffered saline (PBS). Heads were post-fixed in the same fixative for 48 h at room temperature. Brains were harvested and sliced into 50 μm-thick sections using a vibrating microtome (LinearSlicer Pro7, Dosaka EM Co. Ltd, Kyoto, Japan). The formalin-fixed human brain tissues were sliced at 50 μm-thickness and stored in PBS. Dissociated cultures of embryonic (E17~18) mouse hippocampal neurons were prepared from female timed pregnant mice of either non-Tg or P301L-Tg as previously described for rat cultures (Misonou and Trimmer, 2005) with minor modifications. Briefly, dissected hippocampi were digested in 0.25% trypsin for 15 min at 37°C, dissociated by pipetting, and then plated onto glass coverslips coated with 1 mg/ml poly-L-lysine at 50,000 cells/coverslip. Neurons were cultured for 3 to 14 days in vitro (DIV) in 6-well plates, on the bottom of which contained astrocyte cultures. Coverslips were lifted with wax pedestals as described by Kaech and Banker (Kaech and Banker, 2006). Cytosine arabinoflanoside was added to the culture at 2 DIV to prevent the growth of non-neuronal cells on the coverslips. Neurons were fixed with 4% PFA for immunofluorescence staining.

### Immunostaining and visualization of tissue sections and primary cell cultures

The tissue sections and cultured neurons were permeabilized with methanol and blocked in 10% goat serum in PBS with 0.1% Tween 20 (PBS-T) for 60 min. They were then incubated with primary antibodies diluted in 1% BSA-PBS-T for 48 to 72 h at room temperature. After the sections were rinsed with PBS-T, bound antibodies were visualized with secondary antibodies that were conjugated to Alexa Fluor Dyes (Thermo Fisher Scientific, Waltham, MA). The human autopsy tissue sections were stained by Sudan Black B to eliminate lipofuscin autofluorescence (Schnell et al., 1999; Miyasaka et al., 2005b), and then immunostained as above except that the PBS was used in all procedure instead of PBS-T. The specimens were observed under a LSM700 (Carl Zeiss Inc., Jena, Germany), a TCS SP8 LASER scanning confocal (Leica Microsystems, Wetzlar, Germany), or a Leica TCS SP8 STED (Leica Microsystems) microscope using ×10 and ×40 dry objectives and ×63 and x100 oil objectives.

### Microtubule-binding assay

MT-bound and-unbound tau were prepared as described previously with minor modifications (Miyasaka et al., 2010). Briefly, mouse brains were homogenized in MT stabilization buffer (1 mM MgSO_4_, 1 mM EGTA, 0.1 mM DTT, 0.5% Triton X100, 10% glycerol, 20 μM taxol, 2 mM GTP, protease/phosphatase inhibitors, and 0.1 M MES, pH 6.8). After a brief centrifugation, the supernatants were then centrifuged at 135,000 x g for 15 min at 2°C. The levels of tau in the supernatant (MT-unbound fraction) and in the pellet (MT-bound fraction) were quantified by western blotting with purified recombinant tau as standards.

### Quantification of mRNAs and proteins

Total RNA was isolated from the cortices of mouse brains using ISOGEN-LS (Nippon gene, Tokyo, Japan), according to the manufacture’s protocol. Extracted total RNA was subjected to purification of mRNA using OligotexTM –dt30 <Super> mRNA purification kit (Takara). Obtained mRNA was used for the synthesis of first strand cDNA with Takara RNA PCR kit (AMV) (Takara). Subsequently, RT-PCR was performed with cDNA, Power SYBR Green PCR Master Mix (Thermo Fisher Scientific), and primers for the genes of interest (Table 1) on the 7500 Real-Time PCR system (Thermo Fisher Scientific). For the Relative Quantity study, each expression level of mRNA was normalized against mouse GAPDH as an internal control. For the gene chip analysis, total RNA purified from three brains of 1 week or 2 weeks old mice was mixed and processed with the Ambion WT Expression Kit (Thermo Fisher Scientific) according to the manufacturers’ instructions. cRNA was then fragmented, labelled, and hybridized to the Mouse Gene 1.0 ST Arrays using the Gene Chip WT Terminal Labeling and Hybridization Kit (Thermo Fisher Scientific). GeneChip fluidics station 450 was used for processing of the arrays, and fluorescent signals were detected with the GeneChip scanner 3000-7 G. Images were analyzed with the GeneChip operating software (Thermo Fisher Scientific).

**Table 1.**
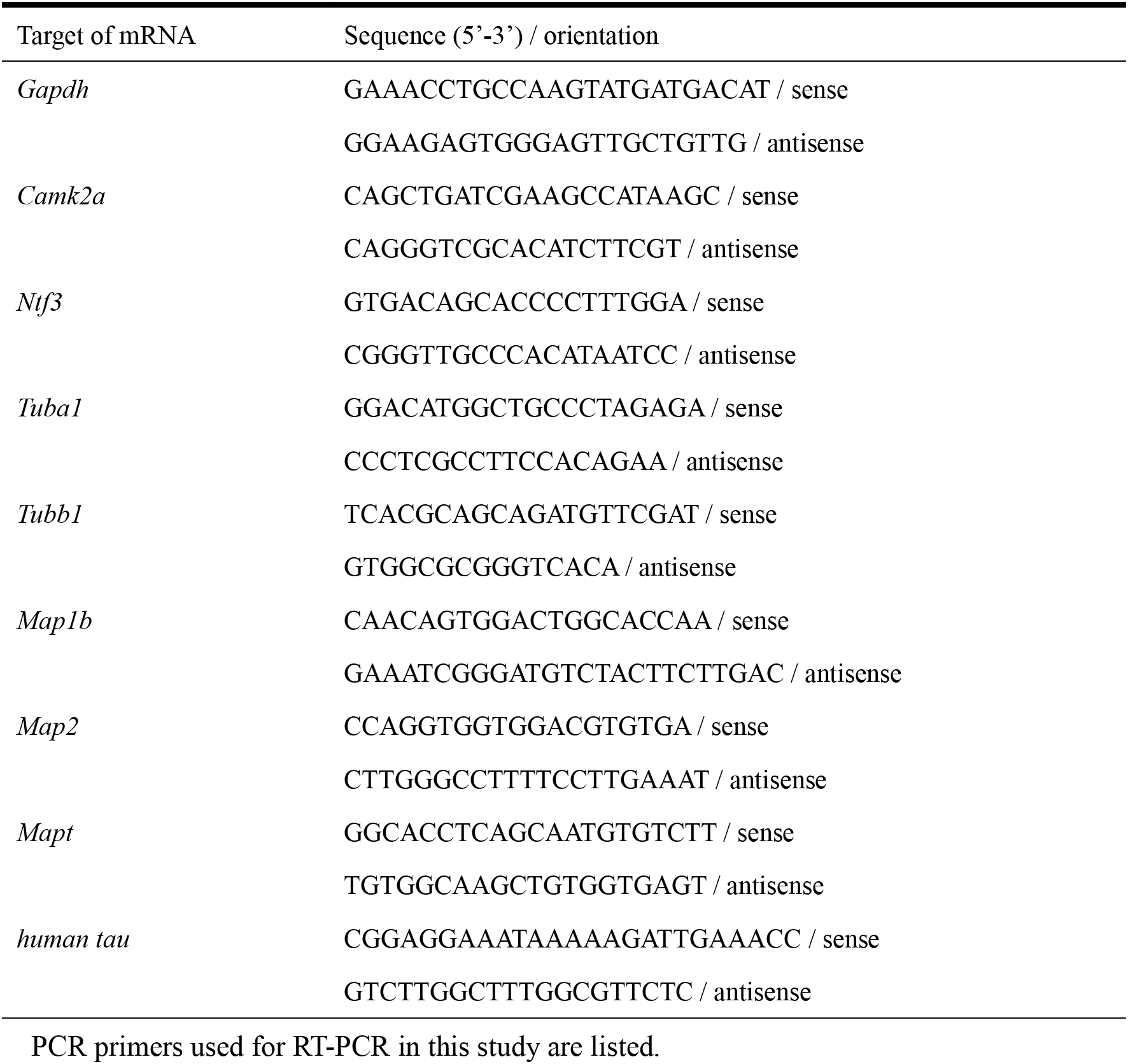
Details of primers used for RT-PCR.

For protein analysis, the right cortical hemisphere was homogenized in O+ buffer (62.5 mM Tris-HCl (pH6.8), 10%(w/v) glycerol, 5% (v/v) 2-mercaptoethanol, 2.3% (w/v) SDS, Protease inhibitors (PMSF, DIFP, pepstatin, Antipain, Aprotinin, leupeptin, TLCK) and Phosphatase inhibitors (NaF, β-glycerophosphate, Na_3_VO_4_, Okadaic acid) at room temperature (Planel et al., 2001). The homogenate was boiled for 5 min and subjected to centrifugation at 100,000 x g for 15 min. The supernatant was solubilized in Laemmli sample buffer and subjected to SDS-PAGE. To assess the levels of proteins, samples were run on 10% acrylamide Tris/glycine gels and subjected to Western blotting with BD-anti-tau for total tau, anti-rodent tauN for mouse tau, tau12 for human tau, or DM1A for α-tubulin. The blots were developed by an enhanced chemiluminescent detection (ECL) (GE HealthCare, Little Chalfont, UK), and the signals were analyzed with a LAS-4000 luminescent image analyzer systems (GE HealthCare). For quantification, the serial diluted samples of wild type mouse brain cortices at 4 weeks old were used as a standard.

### Experimental design and Statistical analysis

Statistical analyses were performed on either GraphPad Prism software (GraphPad Software, Inc). Power analysis was performed using G*Power with parameters taken from previous reports or similar experiments (Faul et al., 2007; Faul et al., 2009).

For the quantification of tau distribution in vivo, we measured fluorescence intensities of mouse (using RTM47) and human tau (with tau12) in the CA3 pyramidal layer and the striatum radiatum, for somatic and axonal signals, respectively, in the P301L-Tg and P301L-KI mouse brain sections. Somatic signals were measured around the nuclei of pyramidal cells, and axonal signals were quantified in the mossy fiber axons. The ratio of somatic and axonal signals were computed, and the averages were analyzed using two-way repeated measures ANOVA with Sidak post-hok test to compare between the ratios of endogenous and exogenous tau in each animal model and between the animal models.

For the assay of tau binding to microtubules, we measured and compared the levels of endogenous mouse tau and exogenous human tau in the unbound fraction in each mouse model using paired Student’s t-test. Also, the levels of human tau in the unbound fraction were compared between the transgenic and knock-in mice using unpaired t-test.

To analyze the expression patterns of endogenous and exogenous proteins in mouse brains, we used regression analyses. We first measured the mRNA levels of tau, tubulins, and CaMKII at different developmental stages. The levels were normalized to the average level at day 1 (for tau and tubulins) or day 70 (for CaMKII) after birth. The normalized levels were plotted against time in days after birth. We then tested if each data set can be fitted with a single exponential function:

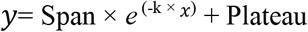

and if a single exponential function with shared slope and plateau can fit the data of tau with either those of tubulins or CaMKII. The latter analysis was supposed to provide us which expression pattern tau follows better. We also compared the span from the exponential curve, which is the difference from the initial value to the plateau value, for each animal. The span therefore reports the direction and magnitude of the change, such that a decrease and an increase in expression would result in positive and negative values, respectively. The mean values were then compared among different mouse proteins using repeated measures ANOVA with the Tukey post-hoc test or between endogenous and exogenous tau in each animal model using paired t-test.

For the protein expression analysis, the ratio of the level at 10 ~ 13 weeks over that at day 1 were computed for both endogenous and exogenous tau in transgenic (P301L-Tg) and knock-in mice. The values were compared using repeated measures Two-way ANOVA with Sidak post-hoc test.

## Results

To investigate the tissue distribution of tau unambiguously, we recently raised anti-tau antibodies, which can distinguish endogenous mouse and exogenous human tau independently of the phosphorylation state of tau, and demonstrated the detailed localization of endogenous mouse tau in non-transgenic (non-Tg) mouse brains (Kubo et al., in press). As shown in Fig. 1., our antibody demonstrates the axon-specific distribution of endogenous mouse tau in the hippocampus, particularly evident in the mossy fiber axons. Using these antibodies, here we investigated how somatodendritic tau emerges in mouse models of tauopathy.

**Figure 1.**
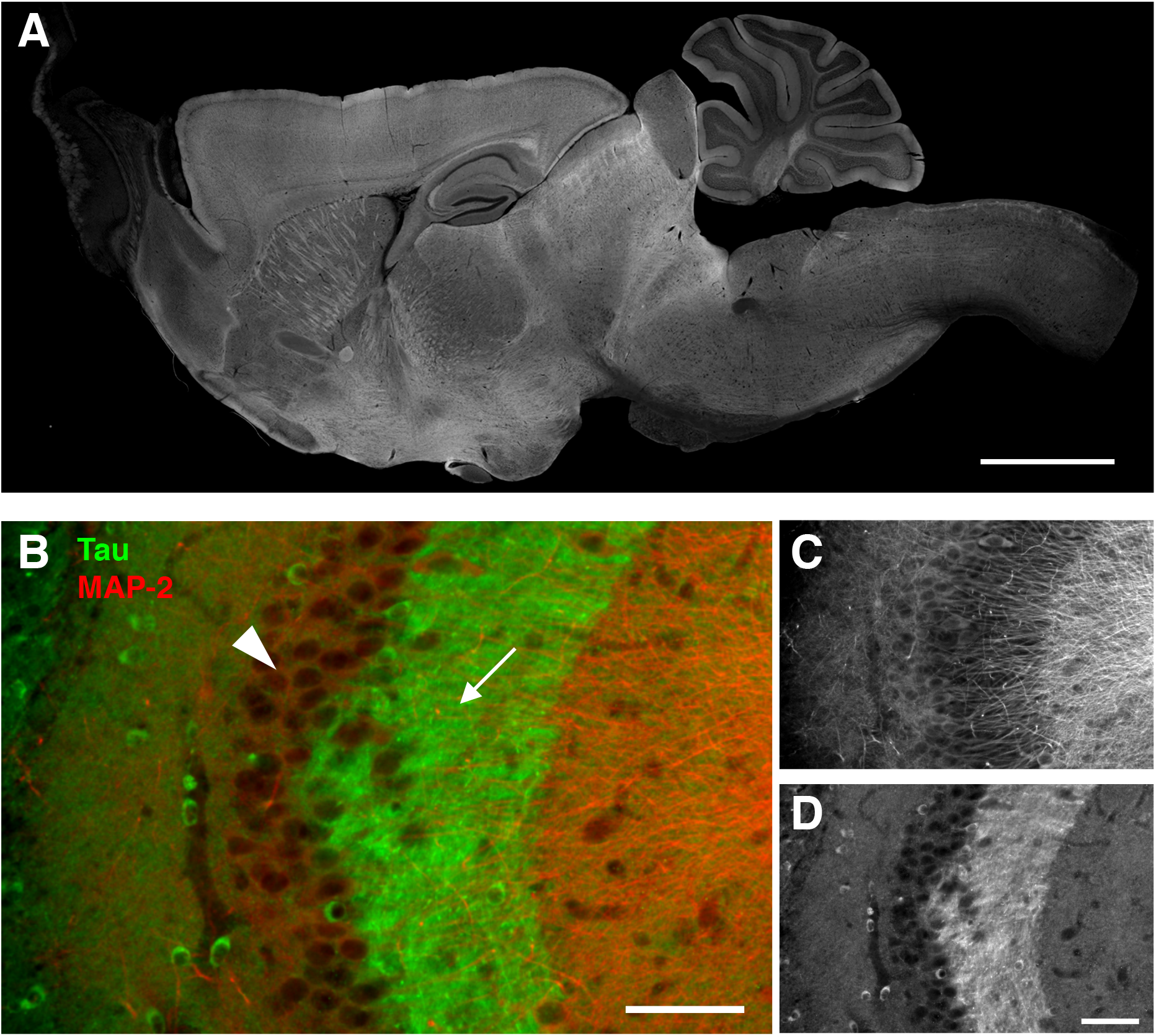
Axonal localization of endogenous tau revealed by our antibodies. ***A*, Whole pictures of the** tissue distribution of endogenous tau in the mouse brain. Sagittal brain sections from perfusion-fixed mice were immunostained with RTM38 anti-tau antibody. Scale bar, 1 mm. ***B - D***, Distribution of endogenous tau in the hippocampal CA3. Tau immunoreactivity (green and ***D***) was abundant in the mossy fiber axons (arrow) but not in the pyramidal layer (arrowhead), where MAP-2 labeling (red and ***C***) is prominent. Scale bars, 100 μm.

We first examined the brains of P301L mutant (P301L-Tg) human tau Tg mice, in which human tau is expressed under the Ca^2+^/calmodulin kinase II (CaMKII) promoter. In this mouse model, neuronal loss and tau pathology occur after the age of 20 months (Kimura et al., 2007; Kimura et al., 2010). At the age of 6 - 12 months, which corresponds to a pre-symptomatic period, endogenous mouse tau, visualized by RTM47 mouse tau-specific antibody, was localized to axons such as mossy fibers in the brains of P301L-Tg (Fig. 2); this pattern was indistinguishable from that in non-Tg mice (Fig. 1). In contrast, exogenous human tau, labeled by human-specific tau12 antibody, was localized not only to the axonal compartment but to the dendrites and cell bodies of CA3 pyramidal neurons (Fig. 2A) as well as in granule cells in the dentate gyrus (Fig. 2B).

**Figure 2.**
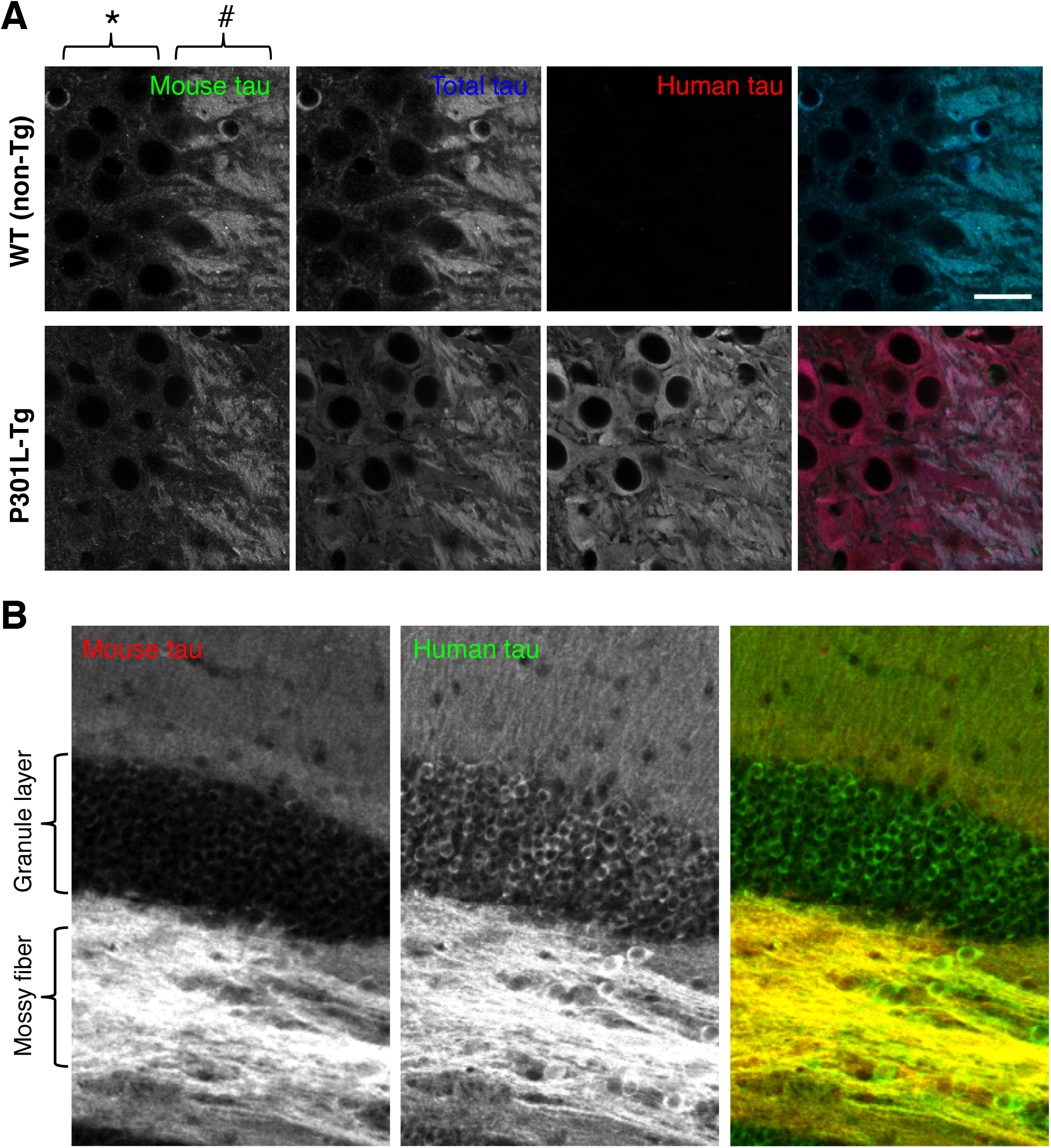
Abnormal somatodendritic localization of exogenous human tau in transgenic mice. ***A***, Hippocampal CA3 area, where we can easily evaluate tau localization in the mossy fiber axons (indicated by #) and adjacent cell bodies of CA3 pyramidal neurons (*), was imaged for endogenous mouse tau, exogenous human tau, and both (total tau). Brain sections from wild type (non-Tg) and P301L mutant Tg (P301L-Tg) mice were labeled with RTM47 (anti-mouse-tau shown in green in composite), anti-tauN (anti-total tau in blue), and tau12 (anti-human-tau in red). The signal intensity of each labeling was adjusted linearly to have a similar intensity in the axons. Scale bar, 20 μm. ***B***, Distributions of mouse (red) and human tau (green) in the dentate gyrus of P301L-Tg mice. Only human tau immunoreactivity was apparent in the granule layer.

The abnormal localization of exogenous tau was not due to the pathogenic FTDP-17 mutation because similar aberrant localization of human tau was observed in the brains of wild-type human tau Tg (WTtau-Tg) mice (Fig. 3). Furthermore, the abnormal localization was not caused by the 3-5 fold greater expression of human tau compared to endogenous tau in these mice (Kimura et al., 2010), as a mild but similar abnormal distribution of exogenous human tau was found in the brains of tau264 mice (Fig. 3), in which the expression level of wild-type human tau is approximately 10% of that of endogenous mouse tau (Umeda et al., 2013). Even at such low expression levels, the exogenous tau was still distributed in cell bodes and dendrites (Fig. 3). This abnormality was also observed in the human-tau expressed in the brains of PS19 mice, a tau-Tg line widely used for the tauopathy research (Fig. 3).

**Figure 3.**
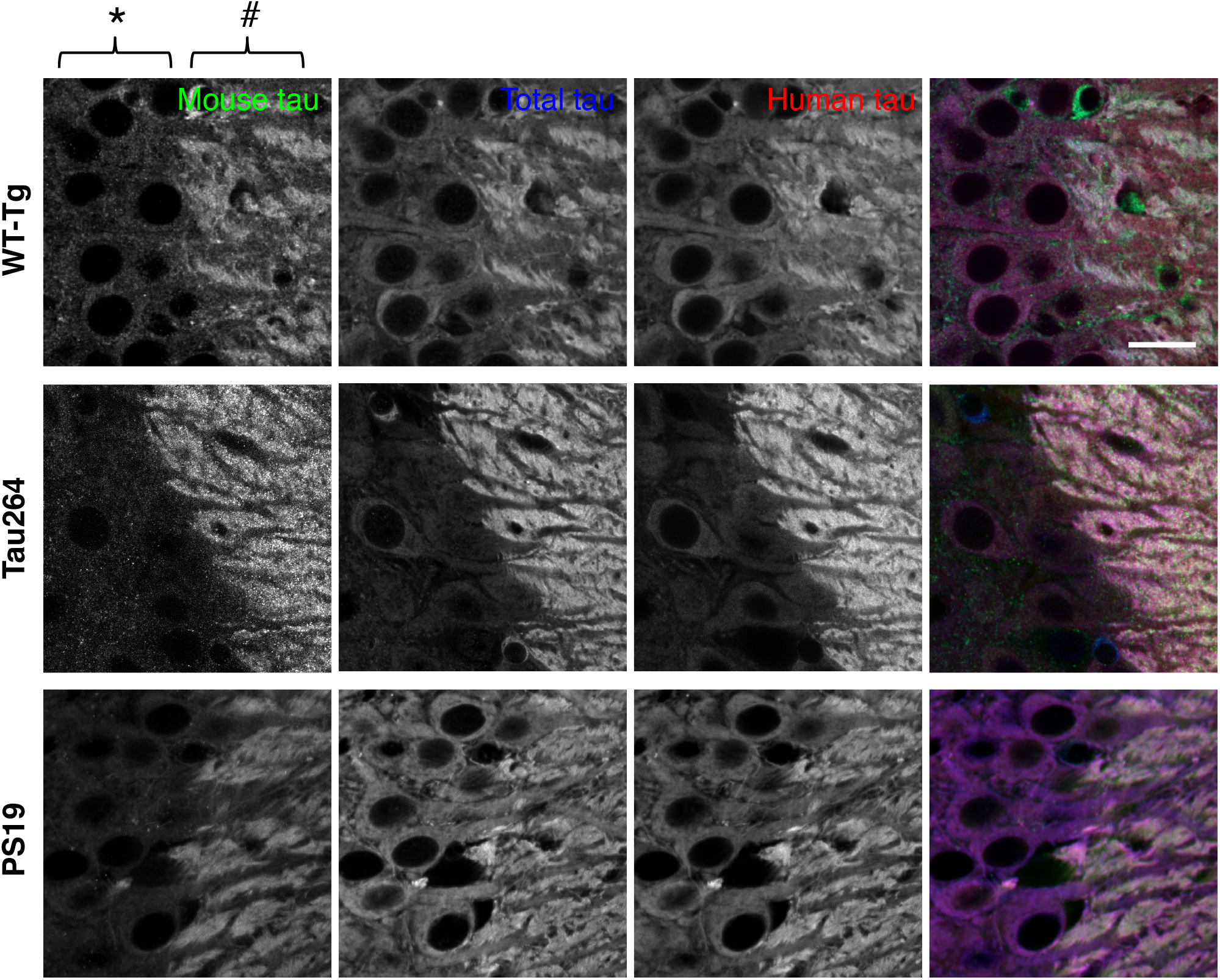
Abnormal somatodendritic localization of exogenous human tau in various transgenic mice. Hippocampal CA3 was labeled for mouse, human, and total tau as in Figure 2A. Transgenic mice expressing high levels of WT human tau (WT-Tg), low levels of human tau (Tau264), and P301S mutant tau (PS19) were used. Mossy fiber axons and adjacent cell bodies of CA3 pyramidal neurons are indicated by # and *, respectively. Scale bar, 20 μm.

The abnormal distribution of exogenous tau could be due to the expression of human tau in mouse neurons. To investigate this cross-species issue, we transfected cultured mouse neurons with mouse tau, of which expression is driven by the CMV promoter. The exogenous GFP-tagged mouse tau appeared to be mis-localized to the soma and dendrites, whereas endogenous mouse tau was properly localized to MAP2-negative axons (Fig. 4), indicating that the mis-localization does not result from the species difference.

**Figure 4.**
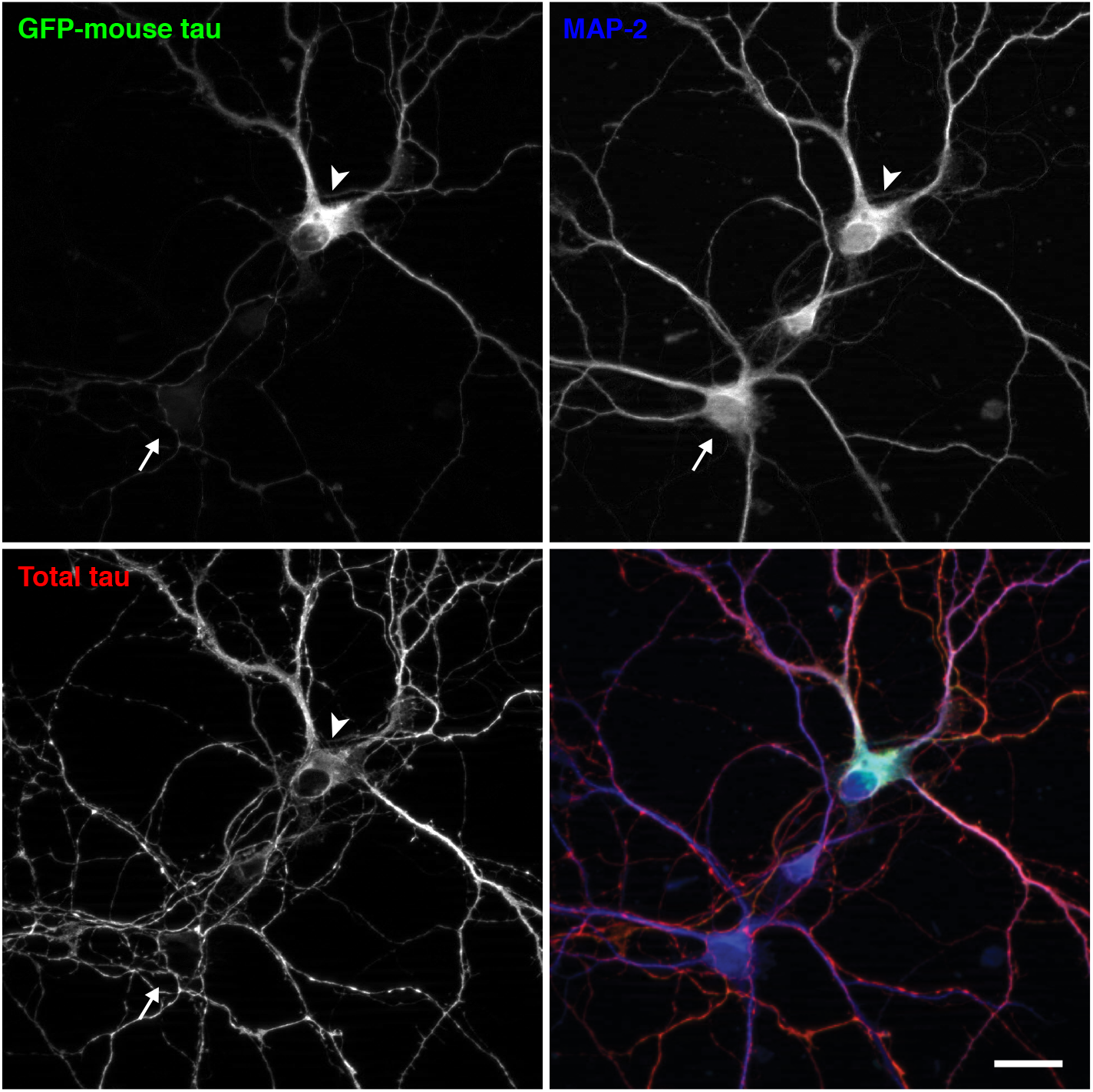
Mis-localization of mouse tau in mouse neurons in culture. Cultured mouse neurons from non-Tg mouse hippocampi were transfected at 7 DIV with EGFP-tagged mouse tau, whose expression is regulated by a CMV promoter. They were fixed at 14 DIV and immunostained for total tau (red in the merged image) and MAP2 (in blue). The arrowhead indicates the soma of a neuron that expresses EGFP-mouse tau, and the arrow indicates the soma of a non-transfected neuron. Scale bar, 20 μm.

We also analyzed the distribution of tau in the brains of P301Ltau knock-in (P301L-KI) mice, in which exogenous human tau replaces endogenous mouse tau and is expressed under the genuine tau promoter. Surprisingly, the exogenous P301L mutant human tau showed a normal axonal distribution, which was particularly apparent in heterozygous mice with endogenous tau (Fig. 5A). The normal axonal localization is not mutation-specific, as V337Mtau also showed axonal localization in V337M-KI mice (Fig. 5B). Line scan analysis and the comparison of axon/soma ratios demonstrated that exogenous tau in KI mice exhibits localization indistinguishable to that of endogenous tau, whereas human tau in P301L-Tg mice shows abnormal somatodendritic localizations (Fig. 5C and D). This is also highly consistent with the finding that these KI mice did not exhibit any tau pathology even at 24 months of age, and even if they were crossbred with APP-Tg mice (Fig. 6). Thus, the abnormal distribution of exogenous tau in dendrite and somata is not due to the species difference nor mutations, but rather the regulatory mechanisms for tau expression.

**Figure 5.**
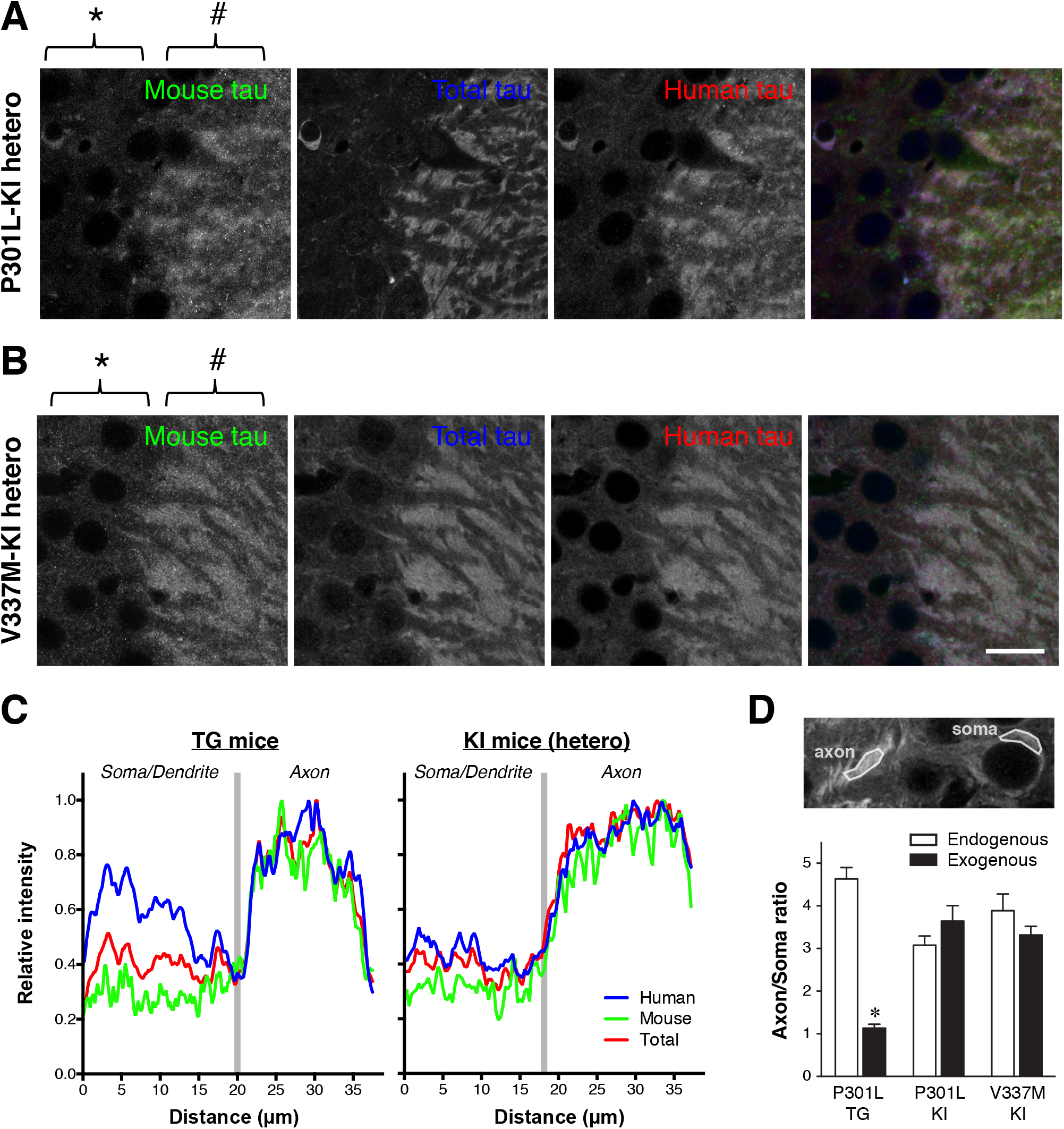
Normal axonal localization of exogenous human tau in P301L tau knock-in mice. ***A***, Mouse and human tau in the hippocampal CA3 of heterozygous and homozygous P301L-KI mice were immunolabeled with anti-mouse tau (RTM47 shown in green), anti-total tau (tauN in blue), and anti-human tau (tau12 in red). It should be noted that, unlike in P301L-Tg mice, both mouse and human tau were virtually absent in the somata (*) but abundant in the axons (#). Scale bar: 20 μm ***B***, Mouse and human tau in the hippocampal CA3 of heterozygous and homozygous V337M-KI mice immunolabeled as in ***A***. Scale bar, 20 μm. ***C***, Normalized intensity profiles of RTM47, anti-tauN, and tau12 in P301L-Tg and V337M-KI mice along lines drawn from the CA3 pyramidal cell layer (*Soma / Dendrite*) to mossy fiber (*Axon*) are shown. ***D***, To quantify the difference in localization between Tg and KI mice, the axon/soma ratio of fluorescence intensity was computed and compared (see *Materials and Methods*). Endogenous mouse tau exhibited high ratios indicating its enrichment in the axon in both mouse models. In contrast, the exogenous human tau in P301L-Tg mice showed significant lower ratios than those of endogenous mouse tau in the same animals (t (10)= 13.87, p < 0.0001 using repeated measures two-way ANOVA and Sidak test) and exogenous human tau in P301L-KI mice (t (20) = 7.012, p < 0.0001) and in V337M-KI mice (t (20) = 5.287, p = 0.0002).

**Figure 6.**
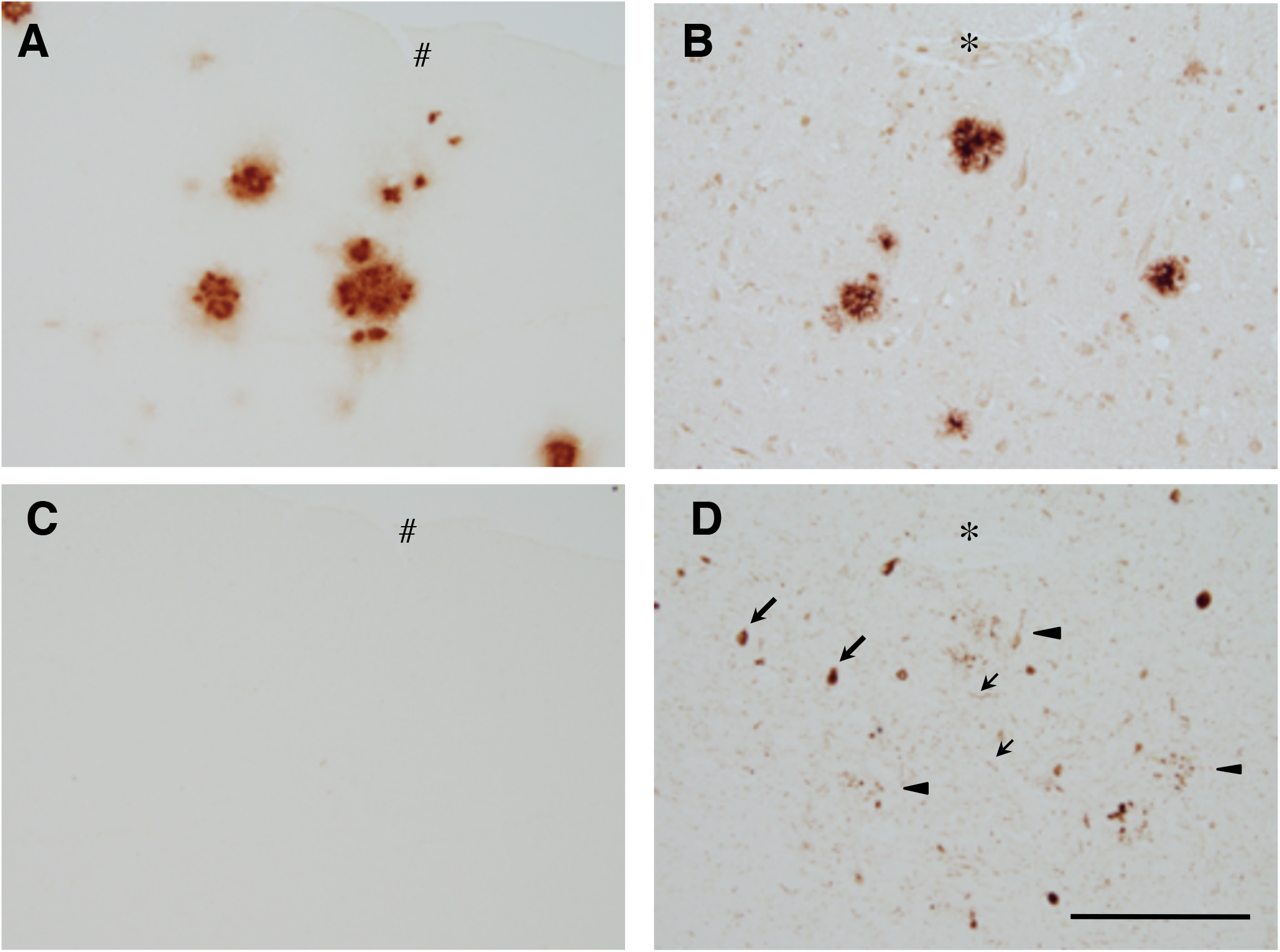
No detectable tau pathology in aged P301L-KI mice. P301L-KI mice were crossbred with APP-Tg mice (Tg-2576), and the amyloid and tau pathologies were examined by immunostaining with anti-amyloid beta and AT8 antibodies at 18 months age (***A*** and ***B***). Sections from AD brain were used for positive control (***C*** and ***D***). Although the abundant senile plaques were detected in both aged crossbred mice and AD brains (***A*** and ***C***), no tau-related pathologies, such as neurofibrillary tangles (large arrows), neuropil threads (small arrows), and dystrophic neurites around senile plaque (arrowheads), were found in the mouse brains (***B***). Asterisks or sharps indicates the same blood vessels appeared on the adjacent brain sections. Scale bar: 100 μm

We recently reported the localization of endogenous and exogenous tau in the Tg mice at super-resolution (at ~50 nm) using the stimulated emission depletion (STED) microscopy (Kubo et al., in press). With the N-terminally binding antibody (anti-tauN), we showed that tau was discontinuously, but not uniformly, distributed in the mossy fiber axons of the hippocampus (Fig. 7A - 7D). These narrow axons with a diameter of 200 - 500 nm are typically packed with several MTs observable with electron microscopy (Amaral et al., 2007). Despite this density of MTs, tau labeling did not appear fibrous but punctate. This unique labeling was reproduced using the RTM38 antibody, of which epitope differs from that of anti-tauN and is in the C-terminal region of tau (Fig. 7E). In contrast, in P301L-Tg mice, exogenous tau in the mossy fiber axons distribute more continuously and appeared fibrous, whereas endogenous mouse tau in the same animals exhibited the punctate labeling (Fig. 8A and B). In contrast, exogenous human tau exhibited the punctate pattern in KI mouse brains (Fig. 8C). These results indicate that the binding of tau to axonal MT is also disrupted in the Tg mice but not in KI mice.

**Figure 7.**
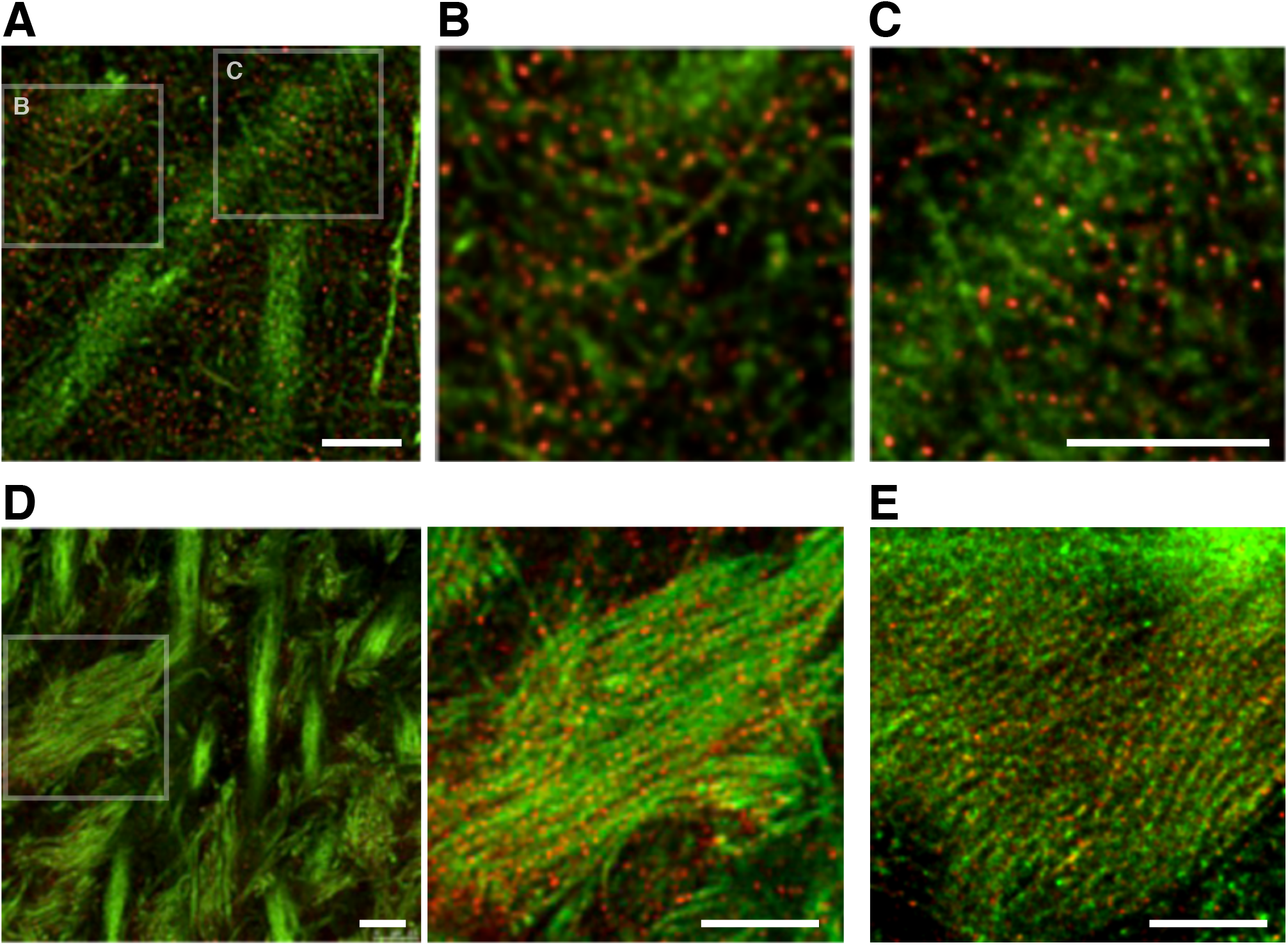
High-resolution imaging of tau in the hippocampal region of non-Tg mouse brains. ***A - C***, Hippocampal areas CA1 in non-Tg mice were labeled with DM1A (green) and anti-tauN (red) and imaged using STED. The boxes in (***A***) indicate the areas that are magnified in (***B***) and (***C***). In CA1, tau is discontinuously localized in axons. ***D***, Hippocampal areas CA3 in non-Tg mice labeled with DM1A (green) and anti-tauN (red). The box in the left image indicate the areas that are magnified in the right image. Punctate tau labeling was observed on the MTs of mossy fibers but was not found in the apical dendrites of pyramidal neurons in area CA3. ***E***, Magnified image of CA3 labeled with DM1A (green) and RTM38 (red). Scale bars: 2.5 μm.

**Figure 8.**
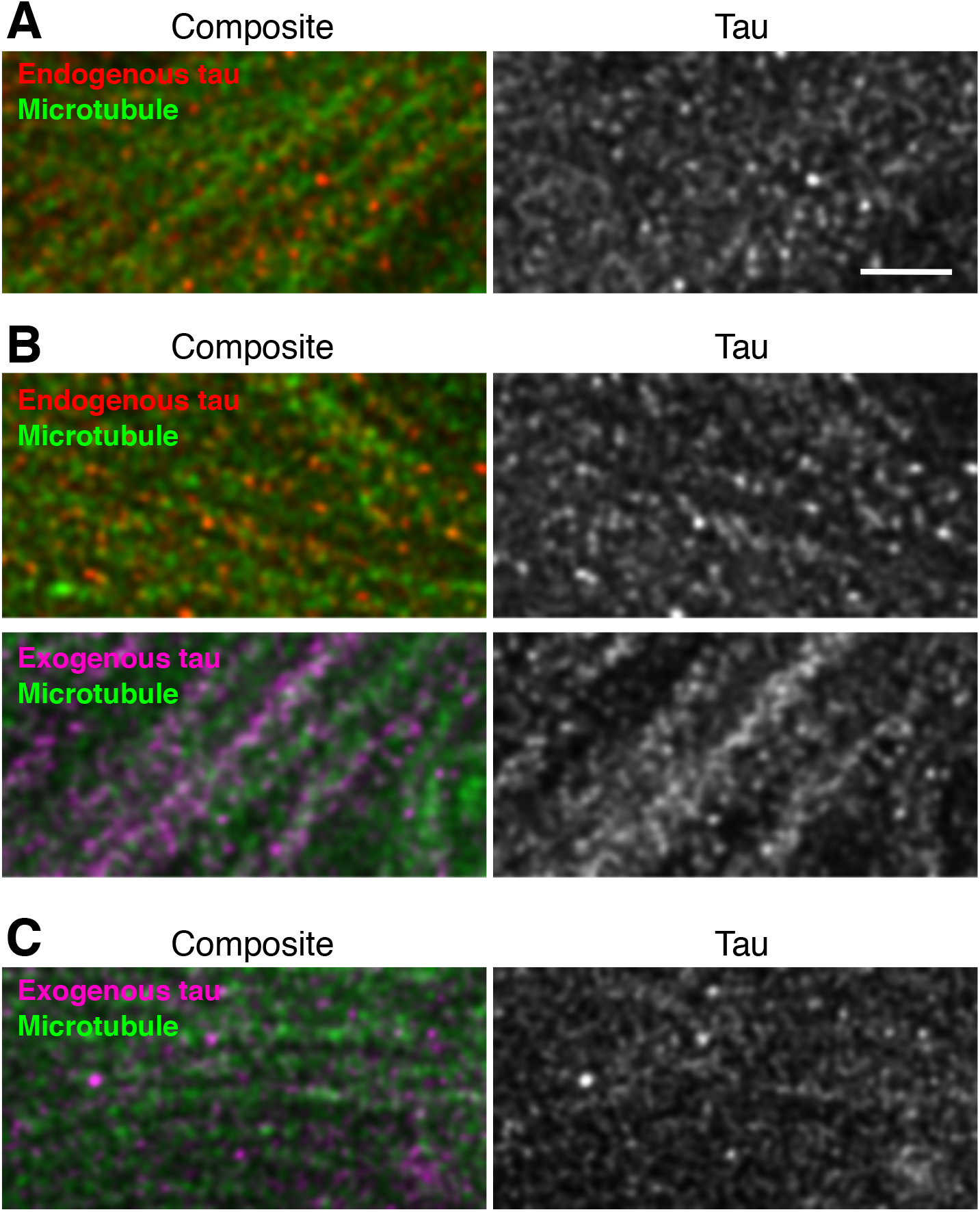
High-resolution imaging of endogenous and exogenous tau in the hippocampi of Tg and KI mouse brains. Mossy fibers of non-Tg (***A***), P301L-Tg (***B***), and V337M-KI (***C***) mice were labeled with anti-RtauN (endogenous tau, red) or RTM49 (human tau, magenta) and observed using STED microscopy. Merged views (left) and tau signal (right) were shown. Endogenous tau exhibited a discontinuous localization pattern, whereas exogenous tau in Tg mice showed a more continuous localization pattern (arrowheads). Scale bar, 1 μm

We therefore examined whether the mis-localization of exogenous tau in tau-Tg mice is associated with abnormal MT-binding biochemically. In the brains of non-Tg mice, the proportion of MT-unbound tau was 23.7 ± 1.7% (Fig. 9A and B). In contrast, in P301L-Tg mice, the proportion of MT-unbound exogenous human tau (64.8 ± 1.4%) was significantly higher than that of MT-unbound endogenous mouse tau (24.9 ± 0.5%) (q (9) = 35.24, *p < 0.0001* using Tukey test after ANOVA, F (2, 9) = 427.7, *p < 0.0001*) or MT-unbound tau in non-Tg mice (q (9) = 36.37, *p < 0.0001*) (Fig. 9A and B). There was no difference in MT-binding of endogenous tau between the non-Tg and P301L-Tg mouse brains (q (9) = 1.128, *p = 0.7137*). Thus, in Tg mice, only exogenous tau is poorly bound to MTs. We then analyzed the MT-binding of exogenous tau in KI mice, in which exogenous mutant tau was localized to the axon in the punctate manner. In these animals, the majority of exogenous tau was found MT-bound, as was the majority of endogenous tau in non-Tg mice (Figure 9C - F). It should be noted that the levels of MT-bound tau depend on how well MTs are preserved during an experiment, such that they differ significantly across experiments. There were no significant differences among the unbound fractions in each model (F (2, 8) = 0.9127, *p = 0.4395* for V337M-KI, and F (2, 9) = 3.879, *p = 0.0610* for P301L-KI using ANOVA). These results suggest that exogenous tau in tau-Ki mice can exhibit a normal axonal distribution and MT-binding if it is expressed under the tau promoter, regardless of mutation.

**Figure 9.**
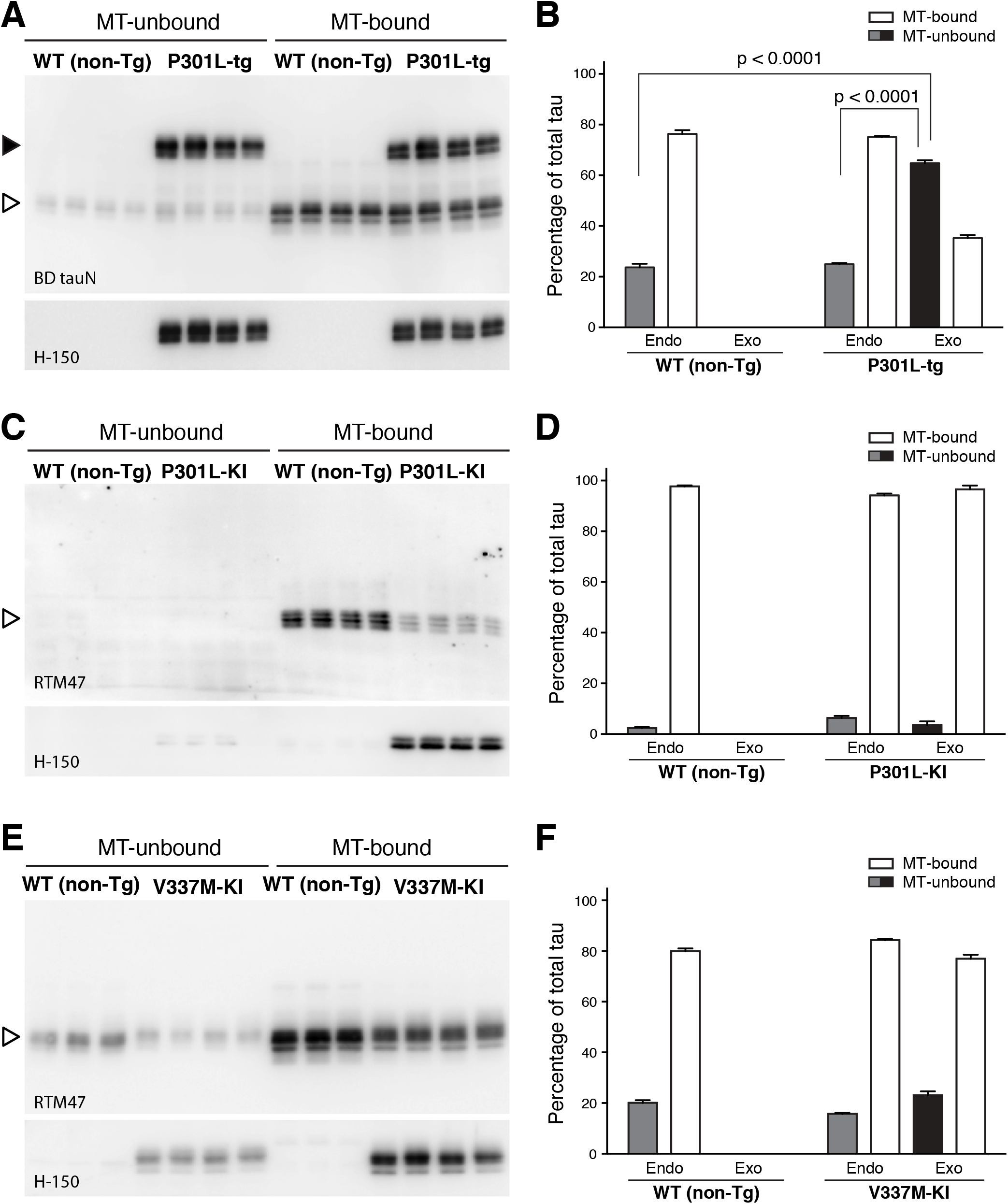
MT binding of human tau in knock-in and Tg mice. MT-bound and-unbound tau were prepared from adult non-Tg and P301L-Tg (***A***), P301L KI (***C***), or V337M-KI mouse brains (***E***) as described in the *Materials and Methods* and detected in Western blotting using anti-total tau (BD anti-tau), anti-human tau (H-150), or anti-mouse tau (RTM47) antibodies. Human and mouse tau in Tg mouse brains are indicated by closed and open arrowheads, respectively. Multiple samples prepared from different animals were shown. Immunoreactive bands of interest were quantified using known amounts of recombinant tau as standards. The proportions of MT-unbound (solid) and-bound (open) endogenous (endo) and exogenous (exo) tau were shown in ***B, D***, and ***F*** for ***A, C***, and ***E***, respectively. The averages of the unbound fraction were compared in each experiment using two-way ANOVA with Sidak post-hoc test because the preservation of MTs and the level of MT-bound tau differ in every experiment. As described in the text, the levels of MT-bound tau vary among experiments due to variable stabilization of MTs. We therefore made comparisons only within each experimental set. The level of unbound human tau was significantly higher than that of endogenous tau in the Tg mice and in WT non-Tg mice (F (2, 9) = 427.7, *p < 0.0001*). In contrast, there was no significant difference in the levels of unbound tau for KI mice (F (2, 8) = 0.9127, *p = 0.4395* for ***D***; F (2, 9) = 3.879, *p = 0.061* for ***F***).

Since our data so far suggest that the ectopic expression of exogenous tau results in the mis-localization as well as abnormal binding to MTs, we investigated how the expression of normal tau and axonal localization occur. First, we investigated the timing when the axonal localization of tau is established during brain development in non-Tg mice. Double-staining with RTM38 anti-tau and anti-MAP2N antibodies showed that tau remains in the cell body and co-localized with MAP2 at day 7 after birth (Fig. 10A). However, at day 14, somatic tau signals were greatly diminished, and tau and MAP2 exhibited discrete distributions in the axon and somatodendrites, respectively (Fig. 10B). Thus, axonal distribution of tau completes between 7 and 14 days after birth. We next examined the distribution of tau in the brains of P301L-Tg mice during developmental stages. While endogenous tau was already localized to the axon, somatodendritic localization of exogenous tau was maintained even at P14 (Fig. 10C). A similar abnormal distribution of exogenous tau was found in cultured hippocampal neurons from embryos obtained by crossbreeding non-Tg and heterozygous P301L-Tg mice. Resulting mixed neurons derived from either non-Tg or Tg embryos could be distinguished based on their immunoreactivity to the human-tau-specific tau12 antibody. In Tg and non-Tg neurons, endogenous tau was detected in the cell body at 3 days in vitro (DIV) (Fig. 10D) but became localized to the axon at 14 DIV (Fig. 10E), as previously reported (Deshpande et al., 2008). Exogenous tau was also found in the cell body at 3 DIV (Fig. 10D), but this somatic expression was maintained even at 14 DIV in Tg neurons (Fig. 10E). Therefore, unlike endogenous tau, exogenous tau, of which expression is driven by the ectopic CaMKII promoter, maintains its somatodendritic localization throughout development and into adulthood.

**Figure 10.**
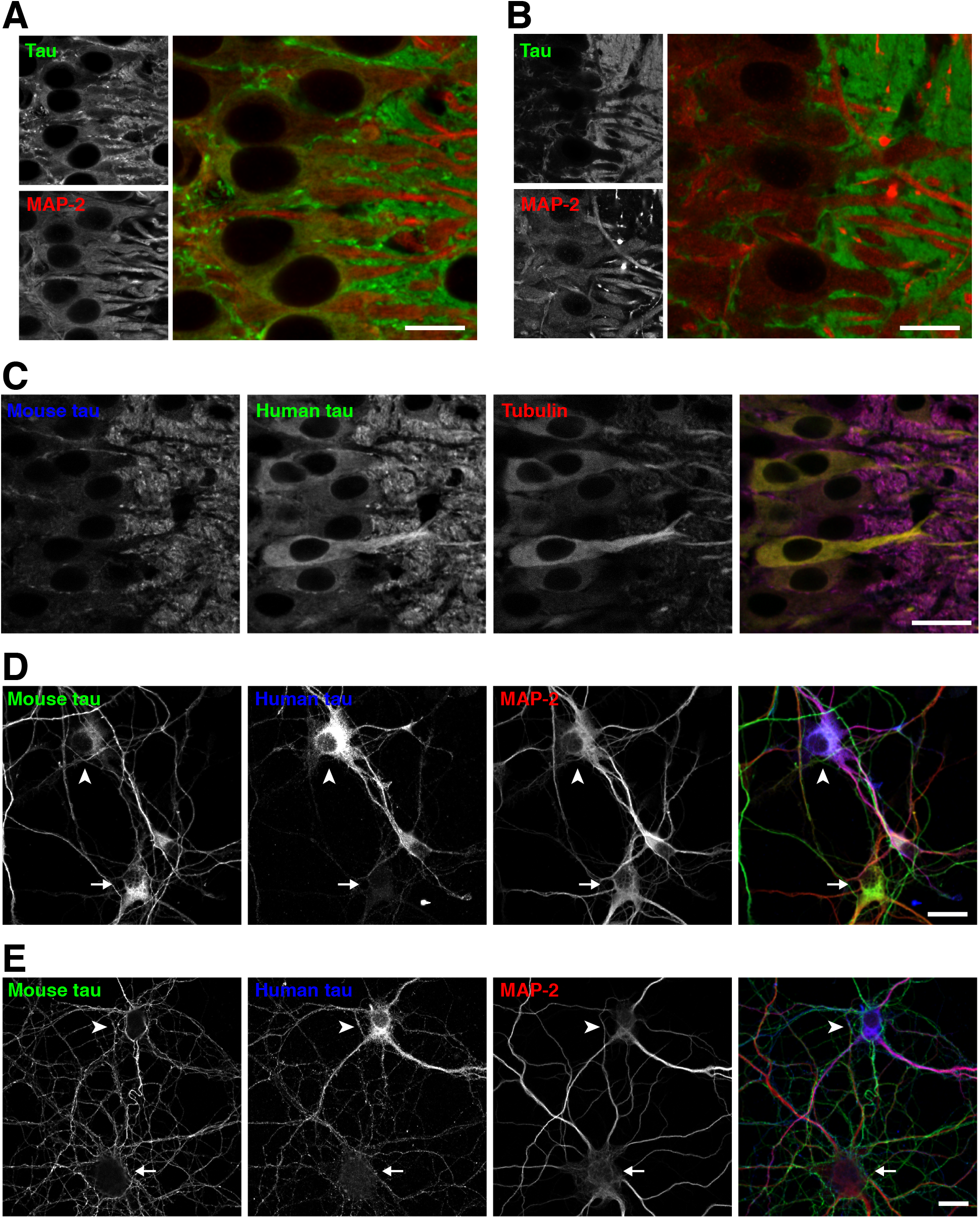
Localization of tau in non-Tg and P301L-Tg mice during development. ***A***, Brain sections from P7 non-Tg mice were subjected to immunostaining for tau (anti-tauN, green) and MAP2 (HM2, red). There were some overlapping signals of tau and MAP-2 in the somata. Scale bar: 20 μm ***B***, Brain sections from P14 non-Tg mice were immunostained for tau (anti-tauN, green) and MAP2 (HM2, red), which exhibited interdigitated patterns at this stage. ***C***, Brain sections from P14 P301L-Tg mice were subjected to immunostaining for mouse tau (anti-RtauN, blue), human tau (tau12, green), and α-tubulin (DM1A, red). Scale bar: 20 μm ***D - E***, Mixed primary cultures of hippocampal neurons from the brains of both non-Tg and P301L-Tg mice were immunostained with anti-mouse tau (RTM47), anti-human tau (tau12), and anti-MAP2N at 3 (***D***) and 14 DIV (***E***). The arrowheads indicate cells that were derived from P301L-Tg mice, and the arrows indicate cells that were derived from non-Tg mice. Scale bars, 20 μm.

Because the promoter of tau expression may dictate its localization in neurons, we investigated how the expression of normal tau is regulated by the genuine tau promoter and how the expression of exogenous tau differs in Tg mouse brains using quantitative real-time PCR. First, the expression of endogenous tau-mRNA (*Mapt*) was quantified in non-Tg mouse cortices at different ages. The level of tau mRNA was highest in the neonatal stage and drastically decreased during the first and second week after birth (Fig. 11A), around the same time that the axonal distribution of tau was completed (Fig. 10A). After 4 weeks, the expression became stabilized at low levels (Fig. 11A). Interestingly, we also found that mRNA levels of α-tubulin (*Tuba1*), β-tubulin (*Tubb1*), MAP-2 (*Map2*), and MAP-1B (*Map1b*) were regulated in a similar fashion (Fig. 11A and Table 2). In contrast, the expression of CaMKII (*Camk2a*), of which promoter was used for the P301L-Tg mouse, exhibited an inverse pattern to that of tau. To analyze and compare these patters quantitatively, we fitted each data set with a single exponential function (see Experimental Procedures). We found that the data of *Mapt*, *Tuba1*, *Tubb1*, *Map2*, and *Map1b* can be fitted well with exponential decay functions, whereas those of *Camk2a* and neurotrophin-3 (*Ntf3*) fit better with exponential growth functions. Particularly, the decay curves for *Mapt*, *Tuba1*, and *Tubb1* had similar parameters, which were statistically indifferent (F (6, 69) = 0.1053, *p= 0.99*). To compare these profiles with that of *Camk2a*, we extracted the span parameter, which reflects the direction and the magnitude of the change in the expression level, from each exponential function (Fig. 11B and Table 2). As expected, spans of *Mapt* and *Tuba1* showed large positive values (1.02 ± 0.16 and 1.06 ± 0.05, respectively), reflecting that they decay over time, whereas the span for *Camk2a* was negative (−1.02 ± 0.04) (Fig. 11C). The overall difference of these values was statistically significant (F (1.202, 3.605) = 168.1, *p= 0.0003*), and there were significant differences between *Mapt* and *Camk2a* (*p = 0.0025*), and *Tuba1* and *Camk2a* (*p < 0.0001*), but not tau and *Tuba1* (*p = 0.95*). The large reduction of tau mRNA expression between 1 and 2 weeks after birth was also confirmed in our gene-chip analysis, such that tau (*Mapt*) was one of the genes which showed more than 2-fold decrease during this period (Fig. 11D). Despite the large changes in mRNA expression during the perinatal development, protein levels of total tau including both 3- and 4 repeat tau were relatively constant (Fig. 11E). These results suggest that the mRNA expression of endogenous tau is highly active only during early development, which presumably results in the constant expression of tau proteins.

**Table 2.** Developmental expression of tau and other genes in the mouse cortex.

| *Gene* | *Span* |
| --- | --- |
| *Tubulins and MAPs* |  |
| *Mapt* | $1.02 \pm 0.16$ |
| *Tuba1* | $1.06 \pm 0.05$ |
| *Tubb1* | $1.08 \pm 0.07$ |
| *Map2* | $0.82 \pm 0.02$ |
| *Map1b* | $0.74 \pm 0.12$ |
| *Other genes* |  |
| *Camk2a* | $-0.95 \pm 0.04^*$ |
| *Ntf3* | $1.05 \pm 0.12$ |

\* $p < 0.0001$ compared to all other genes (F (6, 20)= 66.91).

**Figure 11.**
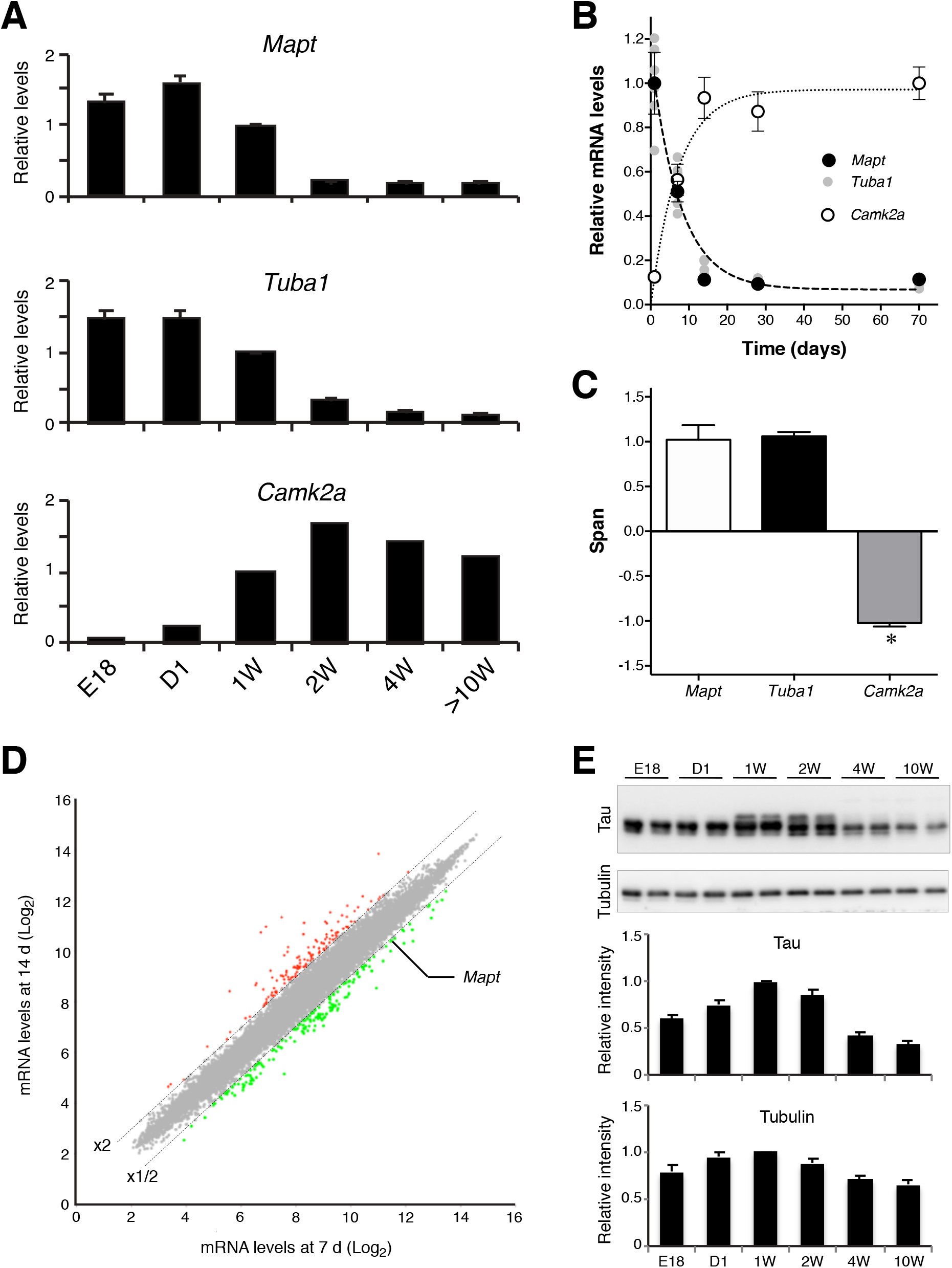
Developmental expression of endogenous tau in the mouse brain. ***A***, Relative levels of mRNA of tau (*Mapt)*, α-tubulin (*Tuba1)*, and CaMKII *(Camk2a)* at the embryonic day 18 (E18), 1 day (D1), 1 week (1W), 2 weeks (2W), 4 weeks (4W), and 10 ~ 13 weeks (10W~) after birth in the cerebral cortices of non-Tg mice were quantified using quantitative real-time PCR. The levels were normalized to that at 1 week for each gene. ***B***, Data in ***A*** were analyzed using nonlinear regression with single exponential curves (upper panel). ***C***, The span of the function in A (see *Materials and Methods*) was obtained from each animal, and the averages were compared among the three genes using repeated-measures ANOVA with Tukey test (F (1.202, 3.605) = 168.1, *p = 0.0003*). ***D***, Microarray analysis comparing cortical transcripts between 1 and 2 weeks after birth. The dashed lines indicate either the 2-fold increase or decrease from the levels at 1 week. Tau (*Mapt*) was identified as one of the genes which exhibited significant decreases. ***E***, Protein levels of tau and α-tubulin. Top, Western blots of tau and tubulin at E18, D1, 1W, 2W, 4W, and >10W using anti-tauN and tubulin antibodies. Duplicates are shown for each time point. Lower panels show the results of quantitation as relative levels to that at 1 week.

The expression profile of *Camk2a* indicated that the expression of human tau in the P301L-Tg mice persists beyond the perinatal period, thereby resulting in the sustained somatodendritic localization of human tau. To test this, we examined the mRNA and protein expression of human tau in this animal model. As shown in Fig. 12A, the level of exogenous tau mRNA in the Tg-mice was lowest at day 1 and continuously increased until adulthood, that is consistent with the *Camk2a* mRNA expression profile (Fig. 12A). In contrast, the expression of tau mRNA in KI mice was at the highest at day 1 and then steeply decreased afterward like endogenous tau (Fig. 12A). We also fitted these data with exponential functions and computed the spans. The results clearly show that the expression of human tau in the Tg mice is inversely regulated compared to that of endogenous tau (Fig. 12B). Concomitantly, the amount of human tau protein in Tg mice also showed a significant age-dependent increase (t (7)= 12.89, p< 0.0001, Two-way ANOVA with Sidak post-hoc test), whereas that in KI mice was relatively constant (t (7)= 2.238, p= 0.12) (Fig. 12C - E). Taken together, these data suggest that excess expression of tau mRNA in mature neurons results in its mis-localization to the somatodendritic compartment and abnormal MT-binding, and may initiate the formation of tau inclusions.

**Figure 12.**
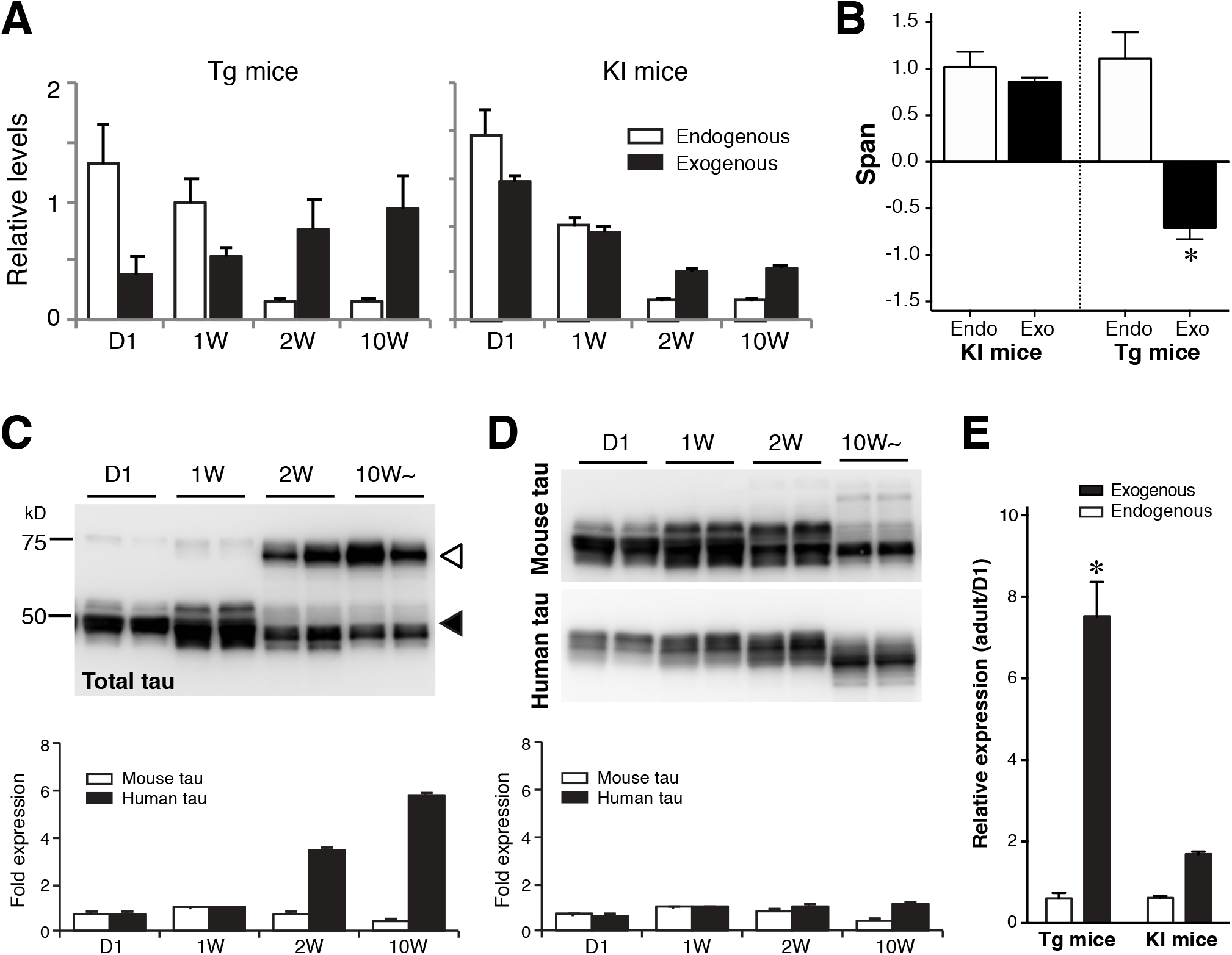
Abnormal expression pattern of exogenous tau in Tg mice. ***A***, Relative expression levels of endogenous mouse tau and exogenous human tau in P301L-Tg and V337M-KI mice. ***B***, Data in ***A*** were fitted with single exponential curves to compute the spans. Average spans were then compared between endogenous and exogenous tau within each model using Two-way ANOVA with Sidak post-hoc test. The difference was significant in the Tg mice (t (6) = 6.13, *p = 0.0017*), but not in the KI mice (t (6) = 0.54, *p = 0.8447*). ***C*** and ***D***, Protein levels of endogenous mouse tau and exogenous human tau in P301L-Tg (C) and KI mice (D) from quantitative Western blotting. The lower panels show relative protein levels to that at 1 week for endogenous (solid) and exogenous (open) proteins individually. ***E***, The ratios of protein levels at D1 and in the adulthood were analyzed for Tg and KI mice. The ratio for exogenous tau in Tg mice was significantly greater than those for endogenous tau and exogenous tau in KI mice (F (1, 14) = 58.81, *p < 0.0001* using repeated measures two-way ANOVA with Tukey tests).

## Discussion

Previous studies of tau focused mostly on abnormal aggregated tau, and those investigating normal tau in vivo have been scarce (Iqbal et al., 2016). The major issue was the lack of immunological reagents sensitive enough to detect non-aggregated tau *in vivo*, as well as those specific for human and mouse tau individually, in vivo. We recently developed several antibodies which can detect human and mouse tau separately at high sensitivity in brain tissues, and showed precise localization of endogenous normal tau in brain tissues (Kubo et al., in press). Using these immunological reagents, we studied how normal tau becomes axonally localized during development in vivo, and what differences make human tau mislocalized in tauopathy mouse models.

The most unexpected result in this study is the dramatic suppression of tau mRNA occurring during the perinatal development. This was particularly interesting because mRNAs of other MT-related proteins (tubulins and MAPs) also showed similar reductions. The timing of mRNA suppression and the timing of axonal localization of tau occur around the same time during the first two weeks after birth. From these findings, we hypothesize that the production of tau protein and its axonal localization occur mostly only during the perinatal development, such that tau expressed beyond this period would mis-localize and exhibit abnormal MT binding. This explains why only P301L-Tg but not P301L-KI mice exhibited the abnormal somatodendritic localization and MT-binding of exogenous human tau.

Tau expression in the Tg mice used here was controlled by the CaMKII promoter, which resulted in the increasing expression of tau mRNA and protein beyond 2 weeks after birth. Intriguingly, this overexpression of exogenous tau did not affect the distribution and MT-binding of endogenous tau, indicating that exogenous human tau proteins do not intermix with endogenous mouse tau proteins within the same neurons. This can only be possible if the majority of abnormal exogenous tau is expressed after the completion of axonal localization and MT binding of endogenous tau, as the hypothesis predicts and our data indicate. The data from the knock-in mice indicate that the regulation of tau expression is mainly achieved at the transcriptional level because the genetic construct used for the tau-KI mice was the human tau cDNA lacking introns and the 3’-untranslated region (UTR) sequence (Sadot et al., 1994).

In the accompanying paper (Iwata et al.), we also demonstrate that mimicking the early and transient expression of exogenous tau in developing neurons, using an inducible expression system, results in the normal axonal localization and colocalization with endogenous tau in culture. In contrast, exogenous tau, which was constitutively expressed, mis-localized to the soma and dendrites in cultured neurons. These results are consistent with the results obtained with Tg- and KI-mice in the present study. It should also be noted that a recent paper reported that endogenous tau edited to have a fluorescent protein also localizes to the axon and significantly differs from overexpressed tau (Xia et al., 2016). Furthermore, when exogenous tau was expressed transiently in relatively mature neurons in culture, it also mis-localized to the soma and dendrites (Iwata et al.). These results also suggest that there is a narrow window during neuronal development, when tau can be efficiently localized to the axon, and that the expression of endogenous tau is precisely orchestrated with this window.

A number of strains of tau-Tg mice that develop tau pathologies in the brains have been reported to date (Noble et al., 2010). Particularly, crossbreeding with an amyloid-producing mouse line or the injection of amyloid β peptide into the brain can cause tau pathology even in Tg mice that express human tau at substantially lower levels (Gotz et al., 2001; Lewis et al., 2001; Umeda et al., 2014). In contrast to these Tg mice, we were unable to find any tau pathologies in the P301L-KI mice, even when they were crossbred with the APP-Tg mice. Therefore, ectopic promoter-driven expression of exogenous tau is critical for the development of tau pathology, at least in rodent brains. Furthermore, the lack of tau pathology in the tau-KI mice suggests that the abnormal distribution of tau in the somatodendritic compartment is a key step for the development of other tau pathologies. Taken together, our study suggests that the dysfunction of the strict transcriptional regulation due to aging or by other factors could be a potential cause of tau mis-localization and abnormal MT binding, which lead to other pathological changes in tauopathies including AD.

Our hypothesis based on the great reduction of tau transcripts after early development requires that the tau proteins generated during the perinatal period is mostly maintained for a long time without significant turnover. This would make tau as one of extremely long-lived proteins in the brain (Toyama and Hetzer, 2013; Toyama et al., 2013). These proteins, such as collagens and histones, are produced mostly during early development and maintained for a long time, sometimes for the lifetime, with very limited turnover. This is consistent with a previous report indicating very slow turnover rates of tau and tubulins in rodent brains (Tashiro et al., 1996). Since ectopically expressed tau behaves abnormally, making tau early and long-lived may be critical for maintaining the physiological function of tau in long axons, particularly unmyelinated axons, where we observed robust tau labeling (Kubo et al., in press).

## Author contributions

A.K. H. Mi Y.I. and T.M. designed the experiments. H.Mi. performed the immunostaining of the primary neuronal cultures. A.N. and M.M. developed the anti-tau rat IgGs. S.W.K. performed the biochemical analyses. A.T., T.T., H.Mo., M.G, and M.I. provided the mouse model. A.K. H.Mi, and T.M. wrote the manuscript.

## Acknowledgements

We thank Akihiro Harada and Fumitaka Oyama for provision of the tau-KI mice. This work was supported in part by the Grant-in-Aid for Scientific Research on Innovative Areas “Brain Protein Aging and Dementia Control” (T.M. 26117004), Challenging Exploratory Research (22650074; T.M.), the “Integrated research on neuropsychiatric disorder”, which was carried out under the Strategic Research Program for Brain Sciences (T.M. and Y.I.), Core-to-Core Program A Advanced Research Networks, the Strategic Research Foundation at Private Universities (S1201009), the Mitsubishi Foundation (T.M.), and the JSPS KAKENHI Grant Number 26640030 (A.K.) and 16K07006 (to H.M.).

## References

Amaral DG, Scharfman HE, Lavenex P (2007) The dentate gyrus: fundamental neuroanatomical organization (dentate gyrus for dummies). Progress in brain research 163:3–22.

Binder LI, Frankfurter A, Rebhun LI (1985) The distribution of tau in the mammalian central nervous system. J Cell Biol 101:1371–1378.

Braak H, Braak E (1994) Morphological criteria for the recognition of Alzheimer’s disease and the distribution pattern of cortical changes related to this disorder. Neurobiol Aging 15:355–356; discussion 379-380.

Braak H, Del Tredici K, Schultz C, Braak E (2000) Vulnerability of select neuronal types to Alzheimer’s disease. Ann N Y Acad Sci 924:53–61.

Dawson HN, Ferreira A, Eyster MV, Ghoshal N, Binder LI, Vitek MP (2001) Inhibition of neuronal maturation in primary hippocampal neurons from tau deficient mice. Journal of cell science 114:1179–1187.

Dehmelt L, Halpain S (2005) The MAP2/Tau family of microtubule-associated proteins. Genome biology 6:204.

Delacourte A, David JP, Sergeant N, Buee L, Wattez A, Vermersch P, Ghozali F, Fallet-Bianco C, Pasquier F, Lebert F, Petit H, Di Menza C (1999) The biochemical pathway of neurofibrillary degeneration in aging and Alzheimer’s disease. Neurology 52:1158–1165.

Deshpande A, Win KM, Busciglio J (2008) Tau isoform expression and regulation in human cortical neurons. FASEB journal: official publication of the Federation of American Societies for Experimental Biology 22:2357–2367.

Faul F, Erdfelder E, Lang AG, Buchner A (2007) G*Power 3: a flexible statistical power analysis program for the social, behavioral, and biomedical sciences. Behavior research methods 39:175–191.

Faul F, Erdfelder E, Buchner A, Lang AG (2009) Statistical power analyses using G*Power 3.1: tests for correlation and regression analyses. Behavior research methods 41:1149–1160.

Feng G, Mellor RH, Bernstein M, Keller-Peck C, Nguyen QT, Wallace M, Nerbonne JM, Lichtman JW, Sanes JR (2000) Imaging neuronal subsets in transgenic mice expressing multiple spectral variants of GFP. Neuron 28:41–51.

Frandemiche ML, De Seranno S, Rush T, Borel E, Elie A, Arnal I, Lante F, Buisson A (2014) Activity-dependent tau protein translocation to excitatory synapse is disrupted by exposure to amyloid-beta oligomers. The Journal of neuroscience: the official journal of the Society for Neuroscience 34:6084–6097.

Ghetti B, Oblak AL, Boeve BF, Johnson KA, Dickerson BC, Goedert M (2015) Invited review: Frontotemporal dementia caused by microtubule-associated protein tau gene (MAPT) mutations: a chameleon for neuropathology and neuroimaging. Neuropathology and applied neurobiology 41:24–46.

Goedert M, Jakes R (1990) Expression of separate isoforms of human tau protein: correlation with the tau pattern in brain and effects on tubulin polymerization. The EMBO journal 9:4225–4230.

Gomez-Isla T, Hollister R, West H, Mui S, Growdon JH, Petersen RC, Parisi JE, Hyman BT (1997) Neuronal loss correlates with but exceeds neurofibrillary tangles in Alzheimer’s disease. Ann Neurol 41:17–24.

Gotz J, Chen F, van Dorpe J, Nitsch RM (2001) Formation of neurofibrillary tangles in P301l tau transgenic mice induced by Abeta 42 fibrils. Science 293:1491–1495.

Harada A, Oguchi K, Okabe S, Kuno J, Terada S, Ohshima T, Sato-Yoshitake R, Takei Y, Noda T, Hirokawa N (1994) Altered microtubule organization in small-calibre axons of mice lacking tau protein. Nature 369:488–491.

Iqbal K, Liu F, Gong CX (2016) Tau and neurodegenerative disease: the story so far. Nature reviews Neurology 12:15–27.

Ittner LM, Ke YD, Delerue F, Bi M, Gladbach A, van Eersel J, Wolfing H, Chieng BC, Christie MJ, Napier IA, Eckert A, Staufenbiel M, Hardeman E, Gotz J (2010) Dendritic function of tau mediates amyloid-beta toxicity in Alzheimer’s disease mouse models. Cell 142:387–397.

Iwata M, Watanabe S, Yamane A, Miyasaka T, Misonou H Companion to this manuscript.

Johnson GV, Jenkins SM (1999) Tau protein in normal and Alzheimer’s disease brain. Journal of Alzheimer’s disease: JAD 1:307–328.

Kaech S, Banker G (2006) Culturing hippocampal neurons. Nature protocols 1:2406–2415.

Kar S, Fan J, Smith MJ, Goedert M, Amos LA (2003) Repeat motifs of tau bind to the insides of microtubules in the absence of taxol. The EMBO journal 22:70–77.

Kimura T, Yamashita S, Fukuda T, Park JM, Murayama M, Mizoroki T, Yoshiike Y, Sahara N, Takashima A (2007) Hyperphosphorylated tau in parahippocampal cortex impairs place learning in aged mice expressing wild-type human tau. The EMBO journal 26:5143–5152.

Kimura T, Fukuda T, Sahara N, Yamashita S, Murayama M, Mizoroki T, Yoshiike Y, Lee B, Sotiropoulos I, Maeda S, Takashima A (2010) Aggregation of detergent-insoluble tau is involved in neuronal loss but not in synaptic loss. The Journal of biological chemistry 285:38692–38699.

Kowall NW, Kosik KS (1987) Axonal disruption and aberrant localization of tau protein characterize the neuropil pathology of Alzheimer’s disease. Ann Neurol 22:639–643.

Kubo A, Misonou H, Matsuyama M, Nomori A, Wada-Kakuda S, Takashima A, Kawata M, Murayama S, Ihara Y, Miyasaka T (2018) Distribution of endogenous normal tau in the mouse brain. J Comp. Neurol. in press

Kuchibhotla KV, Wegmann S, Kopeikina KJ, Hawkes J, Rudinskiy N, Andermann ML, Spires-Jones TL, Bacskai BJ, Hyman BT (2014) Neurofibrillary tangle-bearing neurons are functionally integrated in cortical circuits in vivo. Proceedings of the National Academy of Sciences of the United States of America 111:510–514.

Lewis J, Dickson DW, Lin WL, Chisholm L, Corral A, Jones G, Yen SH, Sahara N, Skipper L, Yager D, Eckman C, Hardy J, Hutton M, McGowan E (2001) Enhanced neurofibrillary degeneration in transgenic mice expressing mutant tau and APP. Science 293:1487–1491.

LoPresti P, Szuchet S, Papasozomenos SC, Zinkowski RP, Binder LI (1995) Functional implications for the microtubule-associated protein tau: localization in oligodendrocytes. Proceedings of the National Academy of Sciences of the United States of America 92:10369–10373.

Matsumura N, Yamazaki T, Ihara Y (1999) Stable expression in Chinese hamster ovary cells of mutated tau genes causing frontotemporal dementia and parkinsonism linked to chromosome 17 (FTDP-17). The American journal of pathology 154:1649–1656.

Misonou H, Trimmer JS (2005) A primary culture system for biochemical analyses of neuronal proteins. Journal of neuroscience methods 144:165–173.

Miyasaka T, Sato S, Tatebayashi Y, Takashima A (2010) Microtubule destruction induces tau liberation and its subsequent phosphorylation. FEBS letters 584:3227–3232.

Miyasaka T, Ding Z, Gengyo-Ando K, Oue M, Yamaguchi H, Mitani S, Ihara Y (2005a) Progressive neurodegeneration in C. elegans model of tauopathy. Neurobiol Dis 20:372–383.

Miyasaka T, Watanabe A, Saito Y, Murayama S, Mann DM, Yamazaki M, Ravid R, Morishima-Kawashima M, Nagashima K, Ihara Y (2005b) Visualization of newly deposited tau in neurofibrillary tangles and neuropil threads. J Neuropathol Exp Neurol 64:665–674.

Noble W, Hanger DP, Gallo JM (2010) Transgenic mouse models of tauopathy in drug discovery. CNS & neurological disorders drug targets 9:403–428.

Planel E, Yasutake K, Fujita SC, Ishiguro K (2001) Inhibition of protein phosphatase 2A overrides tau protein kinase I/glycogen synthase kinase 3 beta and cyclin-dependent kinase 5 inhibition and results in tau hyperphosphorylation in the hippocampus of starved mouse. The Journal of biological chemistry 276:34298–34306.

Planel E, Krishnamurthy P, Miyasaka T, Liu L, Herman M, Kumar A, Bretteville A, Figueroa HY, Yu WH, Whittington RA, Davies P, Takashima A, Nixon RA, Duff KE (2008) Anesthesia-induced hyperphosphorylation detaches 3-repeat tau from microtubules without affecting their stability in vivo. The Journal of neuroscience: the official journal of the Society for Neuroscience 28:12798–12807.

Sadot E, Marx R, Barg J, Behar L, Ginzburg I (1994) Complete sequence of 3’-untranslated region of Tau from rat central nervous system. Implications for mRNA heterogeneity. Journal of molecular biology 241:325–331.

Santacruz K, Lewis J, Spires T, Paulson J, Kotilinek L, Ingelsson M, Guimaraes A, DeTure M, Ramsden M, McGowan E, Forster C, Yue M, Orne J, Janus C, Mariash A, Kuskowski M, Hyman B, Hutton M, Ashe KH (2005) Tau suppression in a neurodegenerative mouse model improves memory function. Science 309:476–481.

Schnell SA, Staines WA, Wessendorf MW (1999) Reduction of lipofuscin-like autofluorescence in fluorescently labeled tissue. J Histochem Cytochem 47:719–730.

Scholz T, Mandelkow E (2014) Transport and diffusion of Tau protein in neurons. Cellular and molecular life sciences: CMLS 71:3139–3150.

Seiberlich V, Bauer NG, Schwarz L, Ffrench-Constant C, Goldbaum O, Richter-Landsberg C (2015) Downregulation of the microtubule associated protein tau impairs process outgrowth and myelin basic protein mRNA transport in oligodendrocytes. Glia 63:1621–1635.

Tashiro T, Sun X, Tsuda M, Komiya Y (1996) Differential axonal transport of soluble and insoluble tau in the rat sciatic nerve. Journal of neurochemistry 67:1566–1574.

Toyama BH, Hetzer MW (2013) Protein homeostasis: live long, won’t prosper. Nature reviews Molecular cell biology 14:55–61.

Toyama BH, Savas JN, Park SK, Harris MS, Ingolia NT, Yates JR, Hetzer MW (2013) Identification of long-lived proteins reveals exceptional stability of essential cellular structures. Cell 154:971–982.

Trojanowski JQ, Schuck T, Schmidt ML, Lee VM (1989) Distribution of tau proteins in the normal human central and peripheral nervous system. J Histochem Cytochem 37:209–215.

Umeda T, Maekawa S, Kimura T, Takashima A, Tomiyama T, Mori H (2014) Neurofibrillary tangle formation by introducing wild-type human tau into APP transgenic mice. Acta neuropathologica 127:685–698.

Umeda T, Yamashita T, Kimura T, Ohnishi K, Takuma H, Ozeki T, Takashima A, Tomiyama T, Mori H (2013) Neurodegenerative disorder FTDP-17-related tau intron 10 +16C--> T mutation increases tau exon 10 splicing and causes tauopathy in transgenic mice. The American journal of pathology 183:211–225.

Viereck C, Tucker RP, Binder LI, Matus A (1988) Phylogenetic conservation of brain microtubule-associated proteins MAP2 and tau. Neuroscience 26:893–904.

Xia D, Gutmann JM, Götz J (2016) Mobility and subcellular localization of endogenous, gene-edited Tau differs from that of over-expressed human wild-type and P301L mutant Tau. Sci Rep, 6:29074.

Xie C, Miyasaka T (2016) The Role of the Carboxyl-Terminal Sequence of Tau and MAP2 in the Pathogenesis of Dementia. Frontiers in molecular neuroscience 9:158.

Xie C, Miyasaka T, Yoshimura S, Hatsuta H, Yoshina S, Kage-Nakadai E, Mitani S, Murayama S, Ihara Y (2014) The homologous carboxyl-terminal domains of microtubule-associated protein 2 and TAU induce neuronal dysfunction and have differential fates in the evolution of neurofibrillary tangles. PloS one 9:e89796.

Yoshiyama Y, Higuchi M, Zhang B, Huang SM, Iwata N, Saido TC, Maeda J, Suhara T, Trojanowski JQ, Lee VM (2007) Synapse loss and microglial activation precede tangles in a P301S tauopathy mouse model. Neuron 53:337–351.

Zempel H, Mandelkow E (2014) Lost after translation: missorting of Tau protein and consequences for Alzheimer disease. Trends in neurosciences 37:721–732.

Zempel H, Thies E, Mandelkow E, Mandelkow EM (2010) Abeta oligomers cause localized Ca(2+) elevation, missorting of endogenous Tau into dendrites, Tau phosphorylation, and destruction of microtubules and spines. The Journal of neuroscience: the official journal of the Society for Neuroscience 30:11938–11950.

